# Proteasome-mediated protein degradation resets the cell division cycle and triggers ESCRT-III-mediated cytokinesis in an archaeon

**DOI:** 10.1101/774273

**Authors:** Gabriel Tarrason Risa, Fredrik Hurtig, Sian Bray, Anne E. Hafner, Lena Harker-Kirschneck, Peter Faull, Colin Davis, Dimitra Papatziamou, Delyan R. Mutavchiev, Catherine Fan, Leticia Meneguello, Andre Arashiro Pulschen, Gautam Dey, Siân Culley, Mairi Kilkenny, Luca Pellegrini, Robertus A. M. de Bruin, Ricardo Henriques, Ambrosius P Snijders, Anđela Šarić, Ann-Christin Lindås, Nick Robinson, Buzz Baum

## Abstract

The archaeon *Sulfolobus acidocaldarius* is a relative of eukaryotes known to progress orderly through its cell division cycle despite lacking obvious CDK/cyclin homologues. Here, in exploring the mechanisms underpinning archaeal cell division cycle control, we show that the proteasome of *S. acidocaldarius,* like its eukaryotic counterpart, regulates the transition from the end of one cell division cycle to the beginning of the next. Further, we identify the archaeal ESCRT-III homologue CdvB as a key target of the proteasome, and show that state-dependent degradation of CdvB triggers archaeal cell division by allowing constriction of a CdvB1:CdvB2 ESCRT-III division ring. These findings suggest an ancient role for proteasome-mediated degradation in resetting the cell division cycle in both archaea and eukaryotes.

---

The cell division cycle is a defining feature of eukaryotic cell biology. It is regulated by a conserved set of proteins found across all eukaryotic phylogenetic groups, indicating that these components were present in the last eukaryotic common ancestor(*1*). Cyclic changes in the activity of cyclin-dependent mitotic kinases order the key transitions in the eukaryotic cell division cycle: CDK-cyclins drive entry into the division cycle, the onset of DNA replication, and entry into mitosis; whereas the loss of CDK-cyclin activity initiates chromosome separation, cell division and origin licensing(*2*). In fact, oscillations in the activity of a single CDK/cyclin pair have been shown to be sufficient to order the cell division cycle in fission yeast(*3*). These observations make the case for cyclin-dependent kinases being the engine of orderly progression through the eukaryotic cell division cycle.

Cyclin-dependent kinases have yet to be identified outside of eukaryotes. However, *Sulfolobus,* a genus of the TACK superfamily of archaea (Thaumarchaeota, Aigarchaeota, Crenarchaeota, and Korarchaeota), possesses an ordered cell division cycle similar in structure to that of many eukaryotes. Within this genus, the established model archaeon *S. acidocaldarius*, exhibits G1 and G2 gap phases that separate discrete rounds of S-phase (DNA replication), and chromosome segregation and cell division(*4*). Significantly, *S. acidocaldarius* cells also possess functional homologues of eukaryotic proteins that execute two of the key cell division cycle events: DNA replication and cell division. Cdc6/Orc proteins mediate replication initiation at multiple origins in both cases(*5*), and the ESCRT-III machinery is responsible for cell division, whereas in eukaryotes it performs a plethora of different membrane processing roles alongside abscission(*6*, *7*). These observations raise a question: in the absence of CDK/cyclins, how do archaeal cells ensure the precise ordering of events as they progress through their cell division cycle?

In the search for proteins that play a common role in the regulation of the cell division cycle in both archaea and eukaryotes, we turned our attention to cell division cycle-dependent protein degradation. In eukaryotes, state-dependent proteasome-mediated degradation of a small number of proteins plays a critical role in the transition from the end of one cell division cycle to the beginning of the next. This is achieved by CDK-driven activation of the ubiquitin-E3 ligase, APC, which triggers proteasome-mediated degradation of cyclin B and Securin, leading to mitotic exit, chromosome segregation(*8*) and cell division(*9*, *10*). While there are no obvious homologues of these proteins in archaea, homologues of the 20S core eukaryotic proteasome are readily identifiable(*11*–*13*). Additionally, the archaeal 20S proteasome appears more conserved in structure and function to the eukaryotic 20S proteasome than any bacterial homologues(*14*). This prompted us to investigate the possibility that proteasome-mediated protein degradation plays a role in the control of cell division cycle transitions in some archaea analogously to that of eukaryotes, albeit without coordination by CDK/cyclins.

With *S. acidocaldarius* as a model TACK archaeon in which to investigate this hypothesis, we begin by studying the structure of its 20S proteasome. We show that the core *S. acidocaldarius* proteasome shares many structural features with the equivalent 20S eukaryotic proteasome, rendering it subject to inhibition by established eukaryotic proteasome inhibitors. Using these inhibitors to test the cellular function of the archaeal proteasome, we go on to show that proteasomal activity is required for these archaeal cells to divide and initiate replication. Proteomics then allowed us to identify a set of proteasomal targets involved in the final stages of the cell division cycle. We identify an archaeal ESCRT-III homologue, CdvB (Saci_1373), which is part of the CdvABC operon (Saci_1374, Saci_1373 and Saci_1372) and previously implicated in *S. acidocaldarius* cell division(*15*–*17*), as a key proteasomal target at the transition from the end of one cycle to the beginning of the next. Strikingly, our analysis suggests that CdvB inhibits cell division by preventing the constriction of another ESCRT-III ring, comprised of the CdvB paralogues CdvB1 (Saci_0451) and CdvB2 (Saci_1416). As a consequence, proteasome-mediated degradation of CdvB triggers cell division in *S. acidocaldarius*.

## Crystal structure of the *S. acidocaldarius* 20S proteasome

To study the function of the 20S proteasome in *S. acidocaldarius*, we carried out a structural analysis to identify potential inhibitors. As shown previously(*18*), recombinant co-expression of the *S. acidocaldarius* 20S core proteasome α- and β-subunits (Saci_0613 and Saci_0662ΔN, respectively) resulted in the formation of an archetypal, but catalytically inactive, twenty-eight subunit 20S proteasomal cylindrical assembly, which we were able to crystallize to determine its three-dimensional structure by X-ray crystallography to a resolution of 3.7 Å (Fig. 1A, Fig. S1, Table S1). Consistent with previously determined archaeal structures(*11*–*13*), this inactive *S. acidocaldarius* core proteasome was found to consist of four stacks of homo-heptameric toroids (with Saci_0662ΔN β-subunits forming the inner rings and Saci_0613 α-subunits forming the outer rings), with a central channel running through it that narrows at the channel entrance to a diameter of 17Å (Fig. 1A). While the 1-14 N-terminal amino-acid residues for all but one of the α-subunits appeared to be disordered and could not be identified in the density map, we were able to model the positions of the missing α-subunit-1 residues 6-14 (AAMGYDR**A**I) in the structure. Interestingly, this N-terminal tail displays a tight turn at Ala13 causing residues 6-12 to point away from the central channel (Fig. S2A). As a result, the “gateway” of the core *S. acidocaldarius* α-ring is present in an open conformation, consistent with our previous functional analyses of the *S. acidocaldarius* Saci_0613/Saci_0662ΔN/Saci_0909ΔN proteasome which, by contrast to the proteasome of eukaryotes(*19*) is proteolytically active in the absence of regulatory ATPases(*18*).

**Figure 1.**
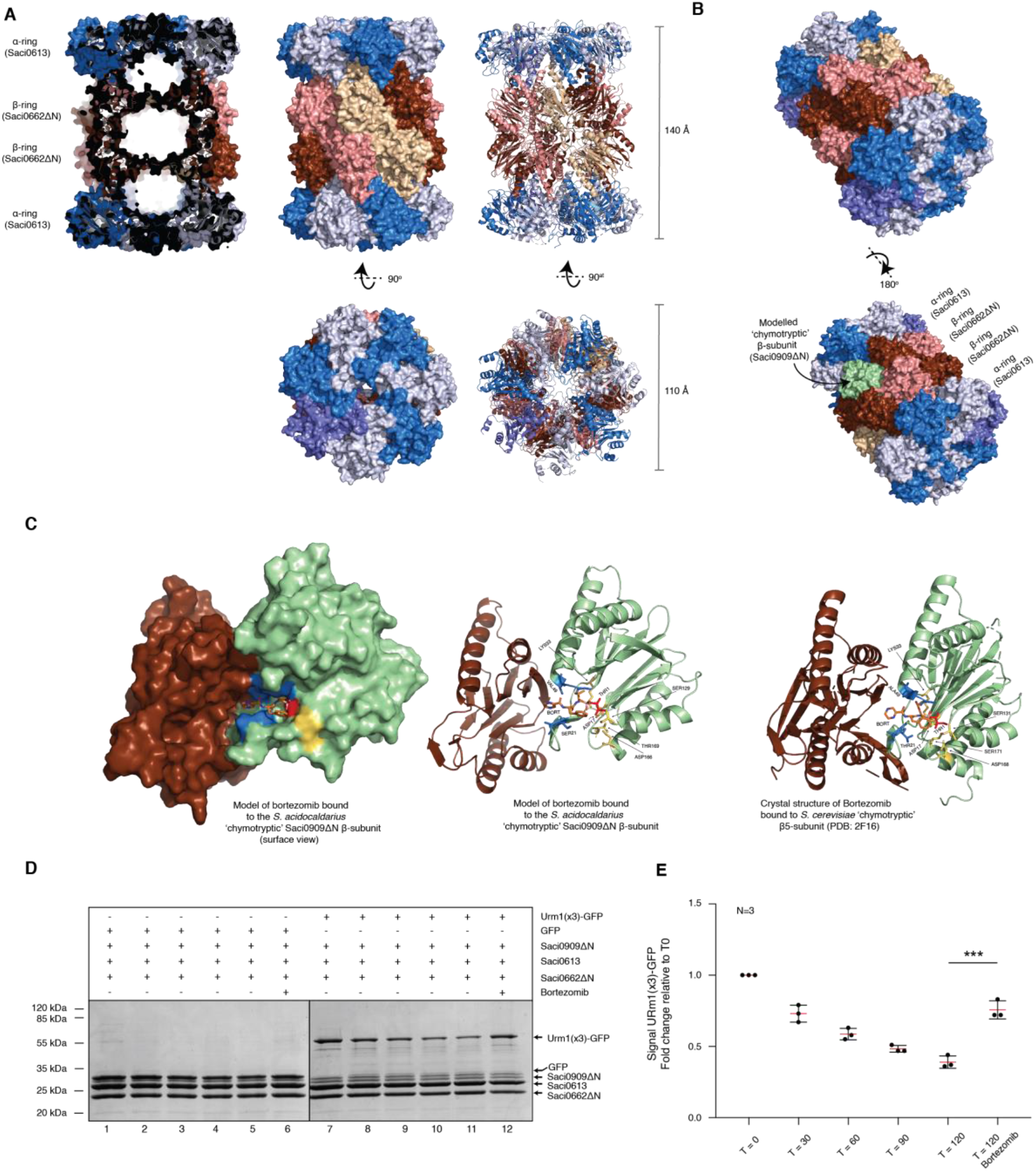
Crystal structure of the *S. acidocaldarius* proteasome. (**A**) (**Top**) Side view of the *S. acidocaldarius* twenty-eight subunit Saci_0613/Saci_0662ΔN 20S proteasome assembly. (**Bottom**) The 20S barrel rotated through 90° to provide a top-down view of the structure highlighting the ‘open’ entrance pores and the translocation channel passing through all three chambers. Saci_0613 α-subunits are coloured slate-blue, and then alternatively dark- and light-blue. Saci_0662ΔN β-subunits are coloured tan, and then alternatively dark- and light-red. (**B**) Modelling of the Saci_0909ΔN catalytic subunit (light green) into the Saci_0613/Saci_0662ΔN 20S crystal structure. (**C**) Modelled binding of the reversible inhibitor bortezomib to the Saci_0909ΔN chymotryptic site in the *S. acidocaldarius* 20S proteasome, using the CovDock software. Surface view (left) and ribbon representation (middle) of the bortezomib inhibitor (orange sticks). The catalytic threonine (Thr1) is shown as a red stick covalently bound to boronic acid group of the bortezomib molecule. The Asp17 and Lys33, Ser129, Asp166 and Thr169, involved in catalysis are shown as yellow sticks. The two loops (blue) harbour the conserved residues Ala20, Ser21 and Gly47 to Val49 mediate binding of the inhibitor. (right) Crystal structure (PDB:2F16) of bortezomib binding at the *S. cerevisiae* 20*S* proteasome focusing on substrate binding pocket formed between the β5 chymotryptic β-subunit (light green) and adjacent β-subunit (brown). (**D**) Time course of a Saci_0613/Saci_0662ΔN/Saci_0909ΔN 20S proteasome assay using a GFP control or a 3xUrm1:GFP substrate. 10 μg of GFP or 3xUrm1:GFP substrates were incubated with 30 μg of the active proteasome complex for 0, 30, 60, 90 or 120 minutes at 63°C (lanes 1-5 and 7-11 for the GFP or 3XUrm1:GFP substrates, respectively). Reactions in lanes 6 and 12 are duplicates of lanes 5 and 11 (GFP and 3xUrm1:GFP incubated with 20S proteasome for 120 minutes at 63°C), but in the presence of bortezomib inhibitor (50 μM). (**E**) Quantification of the Urm1(x3)-GFP signal degradation over time, presented as fold change relative to T = 0 min. N=3. Ratio paired t test, p=0.0006.

As the Saci_0909ΔN β-subunit, which is critical for catalytic activity, was absent from the Saci_0613/Saci_0662ΔN 20S crystal structure, a homology model was generated to determine the likely structure of the biochemically active proteasome(*20*) (see Methods). The final model was determined by its discrete optimized protein energy (DOPE) score(*21*). This structural model of Saci_0909 was then docked into the 20S β-ring in place of one of the seven Saci_0662ΔN β-subunits in the ring (Fig. 1B) using the structures of *Archaeoglobus fulgidus* (PDB ID code 1J2Q) and *Saccharomyces cerevisiae* (PDB ID code 4NNN) as templates for the superimposition.

With the Saci_0909ΔN subunit modelled into the β-ring we were able to predict the structure of the catalytically active sites within the *S. acidocaldarius* 20S core (Fig. S2B), to yield a structure similar to that of its eukaryotic counterpart. Further *in silico* analyses were then performed to assess the likely ability of the small molecule bortezomib (PS-341, Velcade) to inhibit its activity(*22*). A docking model for bortezomib was generated through the CovDock software^23^ (Fig. 1C). This identified a catalytic threonine for covalent attachment to the boronic acid moiety of the inhibitor, as expected(*13*, *23*), and was consistent with the experimentally determined binding state of bortezomib to the *Saccharomyces cerevisiae* 20*S* proteasome (PDB ID: 2F16 and 4QVW)(*24*, *25*) within the β-‘chymotryptic’ site (PDB ID: 2F16)(*24*) (Fig. 1C). Thus, the *S. acidocaldarius* 20S proteasome may process substrates via chymotryptic activity, as suggested for other archaeal and bacterial core proteasomes(*26*).

The model of bortezomib bound to Saci_0909ΔN in the context of the *S. acidocaldarius* 20S proteasome suggested that the small molecule is likely to be an effective inhibitor of its proteolytic activity. Indeed, this compound has been applied as an effective and specific proteasome inhibitor during *in vivo* studies of other archaea(*27*, *28*). As a biochemical test of the inhibitor, we reconstituted the active 20S proteasome assembly (Saci_0909ΔN/ Saci_0662ΔN/ Saci_0613)(*18*) and examined the ability of this complex to degrade a Urm1-tagged GFP substrate in the presence or absence of bortezomib. In this assay, bortezomib effectively inhibited the rate of protein degradation by the active archaeal proteasome (Fig. 1D-E).

## Proteasomal control in archaeal cell division cycle

Having established that it is possible to use the small molecule bortezomib to inhibit the activity of the *S. acidocaldarius* proteasome *in vitro*, we were able to use this to test the function of the archaeal proteasome in cell division cycle control. Given the potential toxicity of long-term inhibition, we restricted treatment to particular stages of the cycle. To achieve this, *S. acidocaldarius* cultures were synchronised to G2 using acetic acid(*29*).

Flow cytometry profiles of cell populations labelled for DNA were obtained and used to characterise the cell division cycle state (Fig. 2A). Following the removal of acetic acid, we observed cells as they progressed through G2, then divided and entered the next division cycle about 80 minutes after release. Strikingly, when cells at this stage in the cycle were treated with the proteasomal inhibitor bortezomib they failed to divide (Fig. 2B-D). This treatment with bortezomib also led to a more than two-fold enrichment (relative to the DMSO control) of cells with compact and separated nucleoids (Fig. 2E-F). The same cell division arrest was observed with MG132, another established eukaryotic proteasomal inhibitor (Fig. S4A).

**Figure 2.**
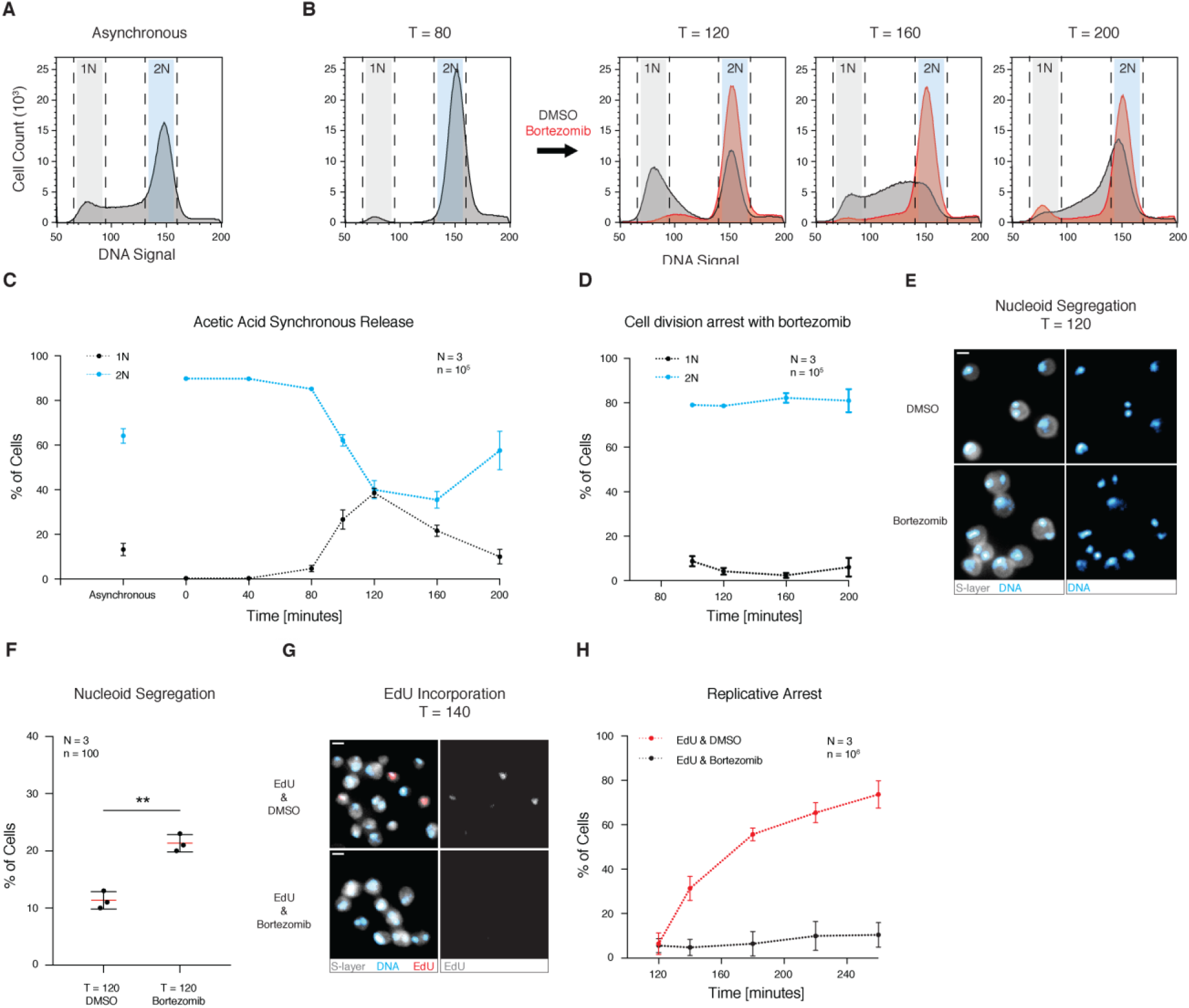
Proteasomal inhibition leads to cell division arrest. (**A**) Histogram shows an asynchronous *S. acidocaldarius* cell population indicating the number of cells with 1N and 2N DNA content. **(B)** Distribution of cells with 1N and 2N DNA content in a population following acetic acid synchronisation and bortezomib treatment 80 min after release (pre-division). N=3, n=10^5^. Representative histograms shown. (**C**) Graph of the distribution of cells with 1N and 2N DNA content in a population following acetic acid synchronisation. (**D**) Graph of the distribution of cells with 1N and 2N DNA content in a population following bortezomib treatment of synchronised pre-division cells 80 min after release. (**E**) Microscopy images of synchronised cells treated with DMSO or bortezomib for 40 minutes, 80 min after release from acetic acid. Fixed cells were stained with Hoechst to visualise DNA content and concanavalin A to visualise the S-layer. Representative images shown. Scale bar = 1 μm. (**F**) Graph of the percentage of cells in the synchronised population with separated nucleoids upon the treatment with DMSO or bortezomib. Ratio paired t-test, p=0.0069. (**G**) Microscopy images of synchronised STK cells treated for 40 min with DMSO or bortezomib, 100 min after release from acetic acid. Fixed STK cells were stained with Hoechst for DNA, concanavalin A for the S-layer, and “click” chemistry was used to label EdU. Scale bar = 1 μm. (**H**) Graph of the percentage of synchronised cells incorporating EdU after DMSO or bortezomib treatment.

To test whether the proteasome inhibitor treated cells were still able to initiate DNA replication while unable to divide, we made use of a *S. acidocaldarius* mutant strain expressing thymidine kinase (STK) generated by the Albers lab(*30*). The introduced thymidine kinase enables *S. acidocaldarius* to incorporate EdU into newly synthesised DNA, which can in turn be observed by “Click-IT” mediated fluorescence microscopy(*30*) (Fig. 2G). The majority of STK cells divided and then incorporated EdU into their DNA as they entered S-phase (before arresting in S-phase (Fig. S3B-C), as reported previously(*30*)). However, EdU was not incorporated into the DNA of pre-division STK cells treated with bortezomib 100 minutes after release from acetic acid arrest (Fig. 2H and Fig. S3A-B). These data show that the activity of the *S. acidocaldarius* proteasome is required for cells to divide and initiate the next round of DNA replication. Importantly, the ability of a proteasome inhibitor to arrest the cycle was stage specific, since the majority of newly divided cells progressed through G1 into S-phase and replicated their DNA on schedule when inhibitors were added to post-division cells 120 minutes after release from the acetic acid arrest (Fig. S3C). Thus, there appears to be a specific role for the proteasome in the regulation of the progression of cells from the end of one cell division cycle to the beginning of the next.

## Proteasomal targets during cell division

Having observed cell division arrest upon proteasomal inhibition (using either bortezomib or MG132), we set out to identify the proteins that are normally subject to proteasome-mediated degradation at division using quantitative mass spectrometry with TMT labelling. The following two conditions were compared: 1) pre-division cells synchronised with acetic acid and arrested with bortezomib and 2) cells collected 15 minutes after the inhibitor was wash-out (Fig. 3A). To understand the role of proteasome-mediated degradation in the cell division process, we focused our attention on the known ESCRT-III-and Vps4-homologous cell division (Cdv) proteins: CdvA (archaea specific), CdvB (ESCRT-III), CdvC (Vps4), CdvB1 (ESCRT-III), and CdvB2 (ESCRT-III). Strikingly, among these functionally-related proteins, CdvB was the only protein significantly and reproducibly enriched in condition (1) compared to condition (2) (Fig. 3A, 3B and Table S2A). In fact, in this experiment CdvB levels proved more sensitive to proteasome inhibition at division than any other protein in the entire proteome (Fig. 3A). A similar enrichment of CdvB was observed when synchronised pre-division cell populations treated with the proteasomal inhibitor MG132 were compared with DMSO treated cells (Fig. S4B, Table S2B). By contrast, CdvA and CdvC were not enriched in bortezomib treated pre-division cells (Fig. 3A), even though they are encoded by the same operon as CdvB, and are part of the same transcriptional wave(*15*, *16*), which remains high in the bortezomib arrest (Fig. S5A). The same was observed for the other two ESCRT-III homologues CdvB1 and CdvB2 implicated in division (Fig. 3A), even though their corresponding mRNAs were expressed at high levels in the pre-division arrest (Fig. S5A). The proteomic data for these proteins were confirmed by western blotting (Fig. 3B, Fig. S5B-C).

**Figure 3.**
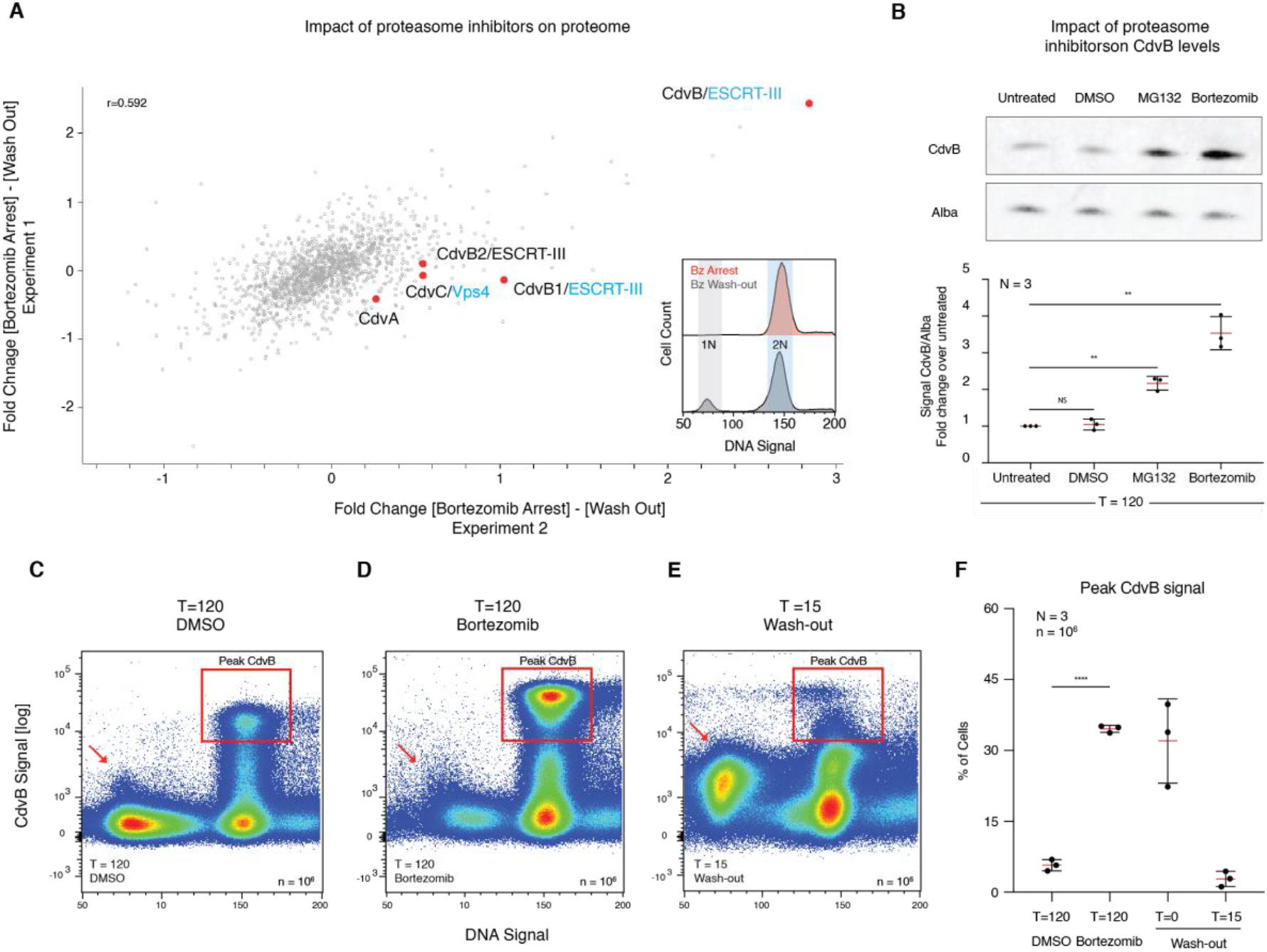
CdvB is the top target of the *S. acidocaldarius* proteasome during cell division. (**A**) Mass spectrometry scatterplot correlating two independent replicates showing the difference in proteomes of (1) pre-division synchronised cells treated with bortezomib for 40 minutes and (2) the population 15 minutes after bortezomib had been washed out. Protein values were measured as the ratio of TMT intensities subtracting sample (2) from sample (1). The insert shows two representative histograms of the bortezomib treated and bortezomib wash out samples as cells state to divide. (**B**) Top, western blots of CdvB and Alba (loading control) for synchronised cells which, 80 min after release from acetic acid, were treated for 40 min with DMSO, MG132, or bortezomib. Bottom, CdvB levels quantified and normalised to Alba, plotted as fold-change relative to the untreated sample. Ratio paired t-test, untreated vs DMSO: p=0.7148, untreated vs MG132: p=0.0041, untreated vs bortezomib: p=0.0033 (**C and D**) 3D scatterplot to show CdvB content vs DNA content vs cell count (colour code: high density red and low density blue) for a synchronised population treated for 40 min with DMSO, and bortezomib, 80 min after release from acetic acid. (**E**) Scatterplot of CdvB content vs DNA content vs cell count, showing a bortezomib-treated synchronised population 15 min after bortezomib has been washed out. (**F**) Quantification of the percentage of cells with peak CdvB fluorescence signal in synchronised populations after DMSO treatment, bortezomib treatment and after washing-out the bortezomib. Unpaired t-test with Welch’s correction, p=<0.0001.

CdvB is part of the ESCRT-III-mediated cell division ring in archaea, and was previously suggested to be a core part of the cell division machinery(*15*–*17*, *31*). Given this, why would an increase in the level of this key division protein be associated with the inability to divide? To explore this question, we used flow cytometry to examine changes in CdvB protein levels in individual cells from synchronised populations as they complete the cell division cycle in the presence or absence of bortezomib, as well as after bortezomib was washed out (Fig. 3C-E). In untreated populations, high levels of CdvB protein were only present in 2N cells and were absent from 1N cells (Fig. 3C). Following treatment with bortezomib, the percentage of cells arrested prior to division with peak CdvB protein content was greatly enriched (Fig. 3D and 3F). Similar results were obtained upon MG132 treatment (Fig. S4C). Then, within 15 minutes of bortezomib wash out, almost all CdvB proteins were degraded as cells progressed through division (Fig. 3F). Taken together, these data suggest that CdvB is normally degraded at a specific time during division in a process that depends on the proteasome.

## Mechanism of archaeal ESCRT-III-dependent cell division

To better understand the role of the proteasome in regulating division itself, we compared the flow cytometry profiles of the archaeal ESCRT-III components, CdvB, CdvB1 and CdvB2 in asynchronous cell populations (Fig. 4A-B, Fig. S6A, and Fig. S7). From these data it is clear that CdvB behaves very differently from CdvB1/B2, in that CdvB is selectively and completely removed from cells during division, while CdvB1/B2 remains at high levels in newly divided cells (Fig. 4A and B, and Fig. S6A). Importantly, when CdvB was plotted against CdvB1 (Fig. 4C) and the DNA content measured as CdvB was degraded (Fig. 4D), it became clear that the bulk of CdvB is degraded just prior to division, with the rest of the protein being lost by the time cells enter G1 (Fig. 4C). Subsequently, levels of CdvB1 and CdvB2 protein gradually decreased to background levels as these cells progressed through S-phase as measured by an increase in the DNA content. Importantly, a similar set of events was seen in flow cytometry profiles in acetic acid synchronised populations as they progressed through division and into the following cycle (Fig. S6B-E).

**Figure 4.**
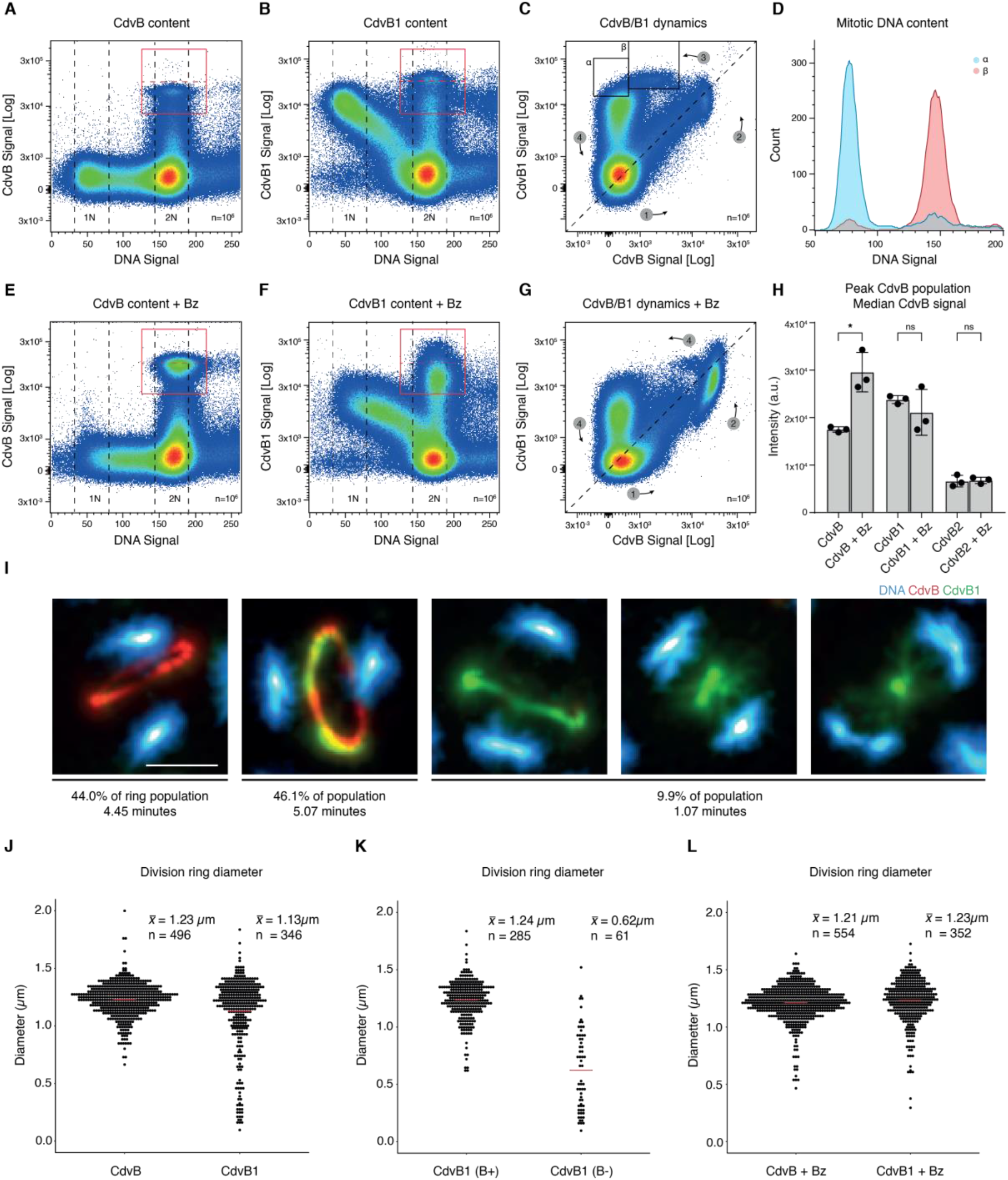
Targeted degradation of CdvB triggers constriction of CdvB1/B2. **(A, B)** Scatterplot of DNA versus CdvB and CdvB1 content for an asynchronous culture. Representative plots shown, N = 3, n = 10^6^. **(C)** Scatterplot of CdvB versus CdvB1 content in an asynchronous culture showing the alternating accumulation and loss of protein (N = 3, n = 10^6^). **(D)** DNA distribution of cells in boxes shown in (C) with ɑ in blue and β in red showing a shift from 2N to 1N. **(E, F)** Scatterplot of DNA versus CdvB and CdvB1 content for an asynchronous culture treated with bortezomib. Representative plots shown, N = 3, n = 10^6^. **(H)** Quantification of CdvB signal in peak populations. Ratio paired two-tailed t test, P = 0.0245. **(G)** Scatterplot of CdvB versus CdvB1 content in an asynchronous culture treated with bortezomib showing an accumulation of cells expressing CdvB and CdvB1 (N = 3, n = 10^6^). **(I)** Representative images of structures observed in asynchronous cultures using SRRF super-resolution microscopy. Fluorescently labelled cells showing Hoechst (blue), CdvB (red) and CdvB1 (green). Percentages indicate total ring bearing population. Minutes of total cell cycle adjusted for exponential age distribution. Scale bar measures 0.5 μm. **(J)** Quantification of CdvB and CdvB1 ring diameters in an asynchronous culture. **(K)** Ring diameters of CdvB1 in the presence (B+) and absence (B−) of a CdvB ring showing how the removal of CdvB leads to a constriction of CdvB1. All quantifications cover more than 6 fields of view over three biological replicates. **(L)** Representative CdvB and CdvB1 rings from cells treated with the protease inhibitor bortezomib showing a loss of constricting population.

In line with this, treating asynchronous populations with bortezomib for 30 minutes led to an increase in the percentage of pre-division cells arrested with peak CdvB and CdvB1/B2 signal (Fig. 4 E-G and Fig. S6F). As expected, treatment with bortezomib was also accompanied by a reduction in the 1N population of cells, as the G1 cells progressed through S-phase (Fig. 4E-F). We also confirmed that whereas bortezomib increased peak protein levels of CdvB per cell, this was not the case for CdvB1/B2 levels (Fig. 4H). Taken together, these findings underlined the selective targeting of CdvB by the proteasome during cell division. To better appreciate how CdvB degradation contributes to the regulation of cell division in *S. acidocaldarius*, we turned to SRRF super-resolution microscopy(*32*). By analysing the structural changes that accompany division, we were able to observe three distinct classes of archaeal ESCRT-III rings: 1) CdvB rings that lack a CdvB1/B2 signal, 2) Rings that contain co-localised CdvB and CdvB1/B2, and 3) CdvB1/B2 rings that lack visible CdvB (Fig. 4I). Cells which displayed CdvB1/B2 but lacked CdvB signal tended to be found in rings of various sizes, as expected if actively constricting. A correlative analysis of CdvB1 and CdvB2 diameters in these images revealed a near perfect correlation between the two proteins (N = 399, r = 0.98, p = 2.2e^−16^) (Fig. S6G). When we analysed the relative sizes of these distinct ring populations, we found that CdvB rings had a reproducible mean diameter of 1.23 μm (N = 496, SD = 0.15) (Fig. 4J). Previous studies have reported a diameter of 0.8 to 1.0 μm for *S. acidocaldarius* cells, and an apparent increase in cell size prior to division(*33*, *34*). In line with this, CdvB1 rings that co-stained for CdvB had a near identical mean diameter of 1.24 μm (N = 285, SD = 0.17). By contrast, CdvB1 rings that lacked CdvB tended to be variable in size and smaller (mean = 0.62 μm, N = 61, SD = 0.35) (Fig. 4K). This supports a role for the selective proteasome-mediated degradation of CdvB from pre-assembled CdvB/CdvB1/B2 rings. Accordingly, we were unable to detect CdvB1 rings that lacked CdvB in cells following treatment with a proteasome inhibitor (Fig. 4L). Further, all of the imaged rings in bortezomib treated cells were found to have a mean diameter equivalent to the width of the CdvB rings seen in the asynchronous populations (mean = 1.23 μm, N = 346, SD = 0.32).

To produce an approximate timeline of events, we used histograms of CdvB vs CdvB1/B2 in exponentially growing asynchronous populations (Fig. 4C) to count the percentage of mitotic pre-division cells. These cells were defined by having 2N DNA content and peak CdvB and/or CdvB1 content. Since the doubling time of these cells is about 3 hours, by adjusting for the exponential age distribution due to binary division(*35*), we found that the average time spent by a cell in a mitotic state was ~11 minutes. Based upon the relative percentages of rings in different states (Fig. 4I), these data suggest that CdvB rings are formed first and have a lifetime of ~260 seconds, before they recruit CdvB1 and CdvB2. The CdvB/B1/B2 rings then persist for an average of ~300 seconds before the proteasome removes CdvB (a process that appears to occur rapidly as we observed few intermediates (Fig. 4C). Finally, CdvB1/B2 rings undergo constriction in a process that lasts ~60 seconds.

## Physical model of ESCRT-III-mediated division in archaea

These findings suggest a simple three-step model for the function of the different ESCRT-III homologues in archaeal cell division. (1) As cells prepare to divide, they assemble a non-contractile CdvB ring of a fixed diameter, recruited via a process that previous studies have suggested is dependent on a CdvA template(*36*, *37*). (2) The initial CdvB ring then acts as a template for the assembly of a CdvB1/B2 polymer which, like force-generating ESCRT-III polymers in other systems, has a smaller preferred curvature(*38*). (3) As a result, the loss of the CdvB scaffold releases potential energy stored in the CdvB1/B2 polymer, driving rapid constriction of the ring as the ESCRT-III-based filaments contract towards their preferred curvature, pulling on the membrane as they do so. The transition between the initial CdvB1(B+)-state and the CdvB1(B−)-state, triggered by the removal of the CdvB-ring, is realized by an instantaneous shortening of the filament bonds in our simulations. Interestingly, when we ran these simulations using various ring radii, we found that the constriction of the ESCRT-III-helix alone was unable to induce division – because of steric hindrance at the midzone (Fig. 5B) (Movie S1, where R/R_0_ = 5%). Thus, only when filaments were allowed to disassemble as they constrict, through the loss of individual subunits from both ends of the helix at a specified rate λ, such that the filament length N(t)=N_0_−λt decreases linearly in time, would cells in the simulations complete cell division (Fig. 5C) (Movie S2, where R/R_0_=5% and λ=2/15t_0_). When we re-analysed data from experiments we observed that while CdvB was not visible in the cytoplasm of cells during division, consistent with its rapid degradation (Fig. 4I), a diffuse CdvB1 signal was seen accumulating as the division ring progressively constricted (Fig. 5E, 5F). The reduction in diameter of the CdvB1/B2 ring was also accompanied by an increase in the local intensity of these proteins in the contracting ring (Fig. 5G). These observations strengthen the conclusions of a previous electron cryo-tomography study by the Jensen and Bell laboratories that revealed both a steady thickening of the contact area between the membrane and an electron dense structure that the authors identified as a putative division ring, and a reduction in ring perimeter during the constriction process(*40*). In summary the experimental observations alongside the physical modelling suggest that ESCRT-III-mediated archaeal cell division requires i) proteasomal degradation of CdvB, and ii) the disassembly of the CdvB1/B2 ESCRT-III polymer as it constricts.

**Figure 5.**
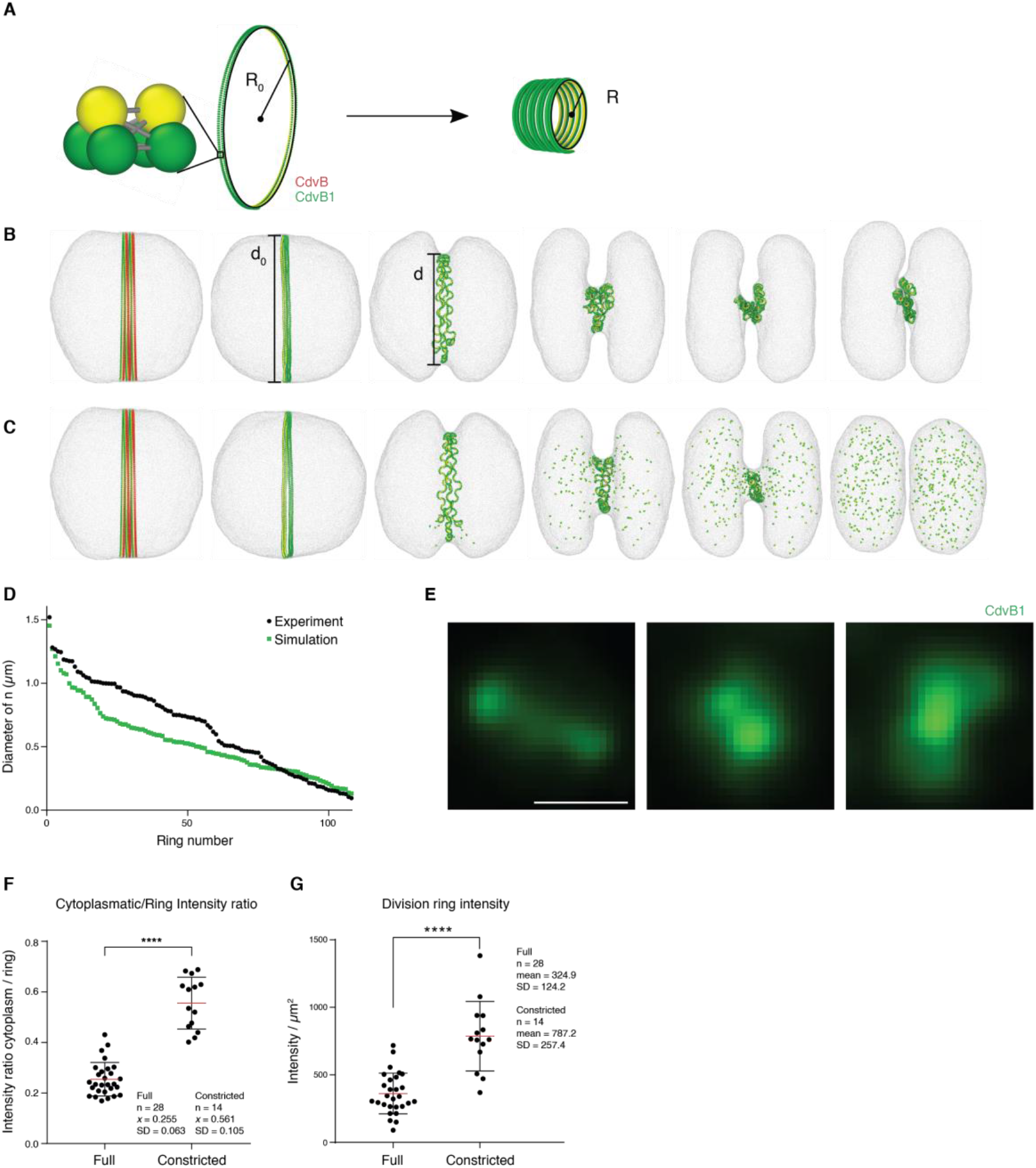
Model of ESCRT-III mediated cell division in archaea. **(A)** The ESCRT filament is built of three-beaded subunits which are connected by harmonic springs to form a helix. Only the green beads are attracted by the cell membrane. The helix undergoes a geometrical change from a state with small local curvature 1/R_0_ (corresponds to CdvB1(B+) to a state with large local curvature 1/R (corresponds to CdvB1(B−). **(B)** Simulation snapshots as a function of time show that cell division is not achieved by constriction of the ESCRT-helix alone; see Supplementary Movie 1. **(C)** Disassembly of the ESCRT-filament is necessary for cell division; see Supplementary Movie 2. **(D)** Progression of division as measured by sampling experimental and simulation data. **(E)** Representative images of CdvB1 showing an increased cytoplasmic signal as constriction progresses. **(F)** Quantification of cytoplasmic signal to ring signal in full and constricted CdvB1 rings. Unpaired Welch’s t test, P = <0.0001. **(G)** Ring signal intensity of full and constricted CdvB1 rings in an asynchronous culture. Unpaired Mann-Whitney test, P = <0.0001.

## Discussion

These findings show how the precise timing of proteasome-mediated protein degradation of a component of the ESCRT-III ring plays a key role in regulating the timing of cell division in *S. acidocaldarius*. This suggests a very simple and elegant mechanism by which *S. acidocaldarius* cells divide: They first assemble a non-contractile ESCRT-III template at the centre of the cell, which is then used to recruit a contractile ESCRT-III ring, whose assembly stores tension in the polymer. This tension is then released once the template is removed and then used to drive rapid cell scission. Further investigation is needed to determine how the ring is positioned, the transitions in states are regulated, and to explore the role of regulatory proteins such as AAA ATPases in targeting CdvB to the proteasome in this process. This study emphasizes the utility of using archaea as a simple model to explore features of molecular cell biology that remain conserved between archaea and eukaryotes. It also suggests the possibility that non-contractile ESCRT-III templates may play an analogous role in the control of ESCRT-III function in some eukaryotic systems. Finally, our study reveals parallels between cell division cycle control in archaea and eukaryotes, implicating proteasome-mediated protein degradation in the regulation of progression from the end of one division cycle to the beginning of the next; in a manner that predates the Cyclin dependent kinases. Further work will be required to identify all the relevant targets of the proteasome during this cell cycle transition in archaea. It will be fascinating to discover whether any of these proteins play common roles in the cell cycle reset in archaea and eukaryotes and/or to determine if there are other shared design features that are used to govern cell cycle progression across the archaeal-eukaryotic divide.

## Supporting information

SMovie_1

SMovie_2

## Acknowledgments

We thank the MRC LMCB at UCL for their support; the flow cytometry STP at the F. Crick Institute for assistance, with special thanks to Sukhveer Purewal and Derek Davis; the members of the Wellcome consortium for archaeal cytoskeleton studies for advice and comments; J. Löwe, S. Oliferenko, M. Balasubramanian and D. Gerlich for discussions and advice on the manuscript. NR and SB would like to thank Neil Rzechorzek, Aline Simon and Salman Anjum for discussion and advice.

## Author contributions

BB and GTR conceived the study. Initial observations were made by FH and GTR. Structural work was planned and carried out by SB and NR, with assistance from DP, MK, and LP. Cell biology methods were developed by GTR, FH, GD, DM and SC with guidance from BB, RH and ACL. Molecular genetics were done by FH and DM. Biochemical analysis was carried out by FH, GTR, AP and LM. RNA analysis was carried out by CF, with guidance from RdB. Mass spectrometry preparation and analysis was carried out by PF, CD, and GTR, with guidance from APS. The physical model was built by AH and LHK, with guidance from AS and BB. The paper was written by GTR, FH and BB, with input from all other authors.

## Data and materials availability

All data is available in the manuscript or the supplementary materials, and raw proteomics data for all validated hits will be made available upon request.

## Supplementary Materials

### Materials and Methods

**Supplementary Figure 1** Electron density map of the Saci_0613/Saci_0662ΔN proteasome

**Supplementary Figure 2** Proteasome open pore and catalytic site

**Supplementary Figure 3** Proteasomal inhibition leads to cell division arrest

**Supplementary Figure 4** Proteasomal inhibition with MG132

**Supplementary Figure 5** Cdv transcript and protein level response to proteasomal inhibition

**Supplementary Figure 6** Targeted degradation of CdvB triggers constriction of CdvB1/B2

**Supplementary Figure 7** In vitro analysis of CdvB, CdvB1 and CdvB2 antibody specificities

**Supplementary Table 1** Data Collection and refinement statistics

**Supplementary Table 2** Mass Spectrometry tables with Cdv proteins

**Supplementary Table 3** List of antibody host species and secondary antibodies used

**Supplementary Table 4** List of primers used in cloning of plasmids used for protein expression

**Supplementary Table 5** List of primers used for RT-qPCR analysis

**Supplementary Movie 1** Constriction simulation without filament disassembly

**Supplementary Movie 2** Constriction simulation with filament disassembly

**References** (*41*–*52*)

