## Supplementary Information for "Proteasome-mediated protein degradation resets the cell division cycle and triggers ESCRT-III-mediated cytokinesis in an archaeon"

**This PDF file includes:**

Materials and Methods
Figs. S1 to S7
Tables S1 to S5
Captions for Movies S1 to S2

**Other Supplementary Materials for this manuscript include the following:**

Movies S1 to S2

#### Materials and Methods

##### Strains, culture media and growth conditions

The *S. acidocaldarius* strains DSM639, MW001 and STK, a thymidine kinase mutant(*30*), were grown at 75°C in Brock medium supplemented with 0.1% NZ-amine and 0.2% Sucrose[33]. The density of liquid cell cultures was maintained at OD600 values between 0.050 and 0.300 as measured by a spectrophotometer. This optical density corresponds to the known exponential growth phase of *S. acidocaldarius.*

##### Synchronisation and drug administration

*S. acidocaldarius* was arrested by treating a population with 3 mM acetic acid for 4.5 hours, approximately 1.5 times the doubling time(*29*). The proteasomal inhibitor bortezomib was used at a final concentration of 10uM, EdU to a final concentration of 200 µM, and MG132 to a final concentration of 100uM. All drugs were dissolved in DMSO.

##### Protein extraction

25 ml *S. acidocaldarius* pellets were lysed in 100-250 µl lysis buffer (TK150 buffer, supplemented with DNAse1, and EDTA-free protease inhibitor cocktail, 0.1% Triton X-100). Cells were disrupted in a cup sonicator. After clarification, protein concentration was measured using a Bradford assay (Bio-Rad).

##### RNA extraction

TRIzol Reagent (Invitrogen) was used to isolate total RNA from 25 mL *S. acidocaldarius* pellets. The ThermoFisher trizol-reagent protocol was used for extraction. A sample clean-up was used to remove residual contaminants in which 3M $Na_4OAc$ (1:10) and 100% ethanol (5:2) was added following RNA extraction. Samples were then incubated overnight at -20°C before washed with 75% ethanol, and spun down at 12,000 rcf at 4°C for 5 minutes. Samples were resuspended in 100µl autoclaved double-distilled water. RNA concentration and purity was measured by NanoDrop spectrophotometer. 260/280 and 260/230 absorbance ratios were approximately 2.0.

##### Quantitative reverse transcription PCR (RT-qPCR)

The abundance of mRNA for selected genes was measured by RT-qPCR. For each reaction, 80 ng of RNA was combined with a Euroscript reverse transcriptase (Eurogentec), 2x MESA BLUE PCR reaction mix (Eurogentec) and the primer pair (Table S5) for each gene, to reach a final volume of 14 µl. Each reaction was run in triplicate and each experiment was run in three biological replicates.

RT-qPCR was run on a Bio-Rad CFX Connect Real-Time System. Each run has 39 cycles of denaturation at 95°C for 3 seconds, annealing and extension of primers at 60°C for 45 seconds, followed by measurement of fluorescence intensity. The relative fold difference between RNA abundance of each gene was derived using the $2^{-\Delta\Delta C(T)}$ method for data normalisation, with equations:

$$\Delta C(T)= C(T)_{\_GENE} - C(T)_{\_REFERENCE\ GENE}$$

$$FD=2^{(-\Delta C(T))}$$

C(T) indicates the number of PCR cycles required to reach a threshold in fluorescence intensity of the sample. C(T) is collected for the gene of interest and reference housekeeping gene, SecY (Saci_0574) as the average of three replicates.

##### ELISA

Antibody specificity was measured using 96-well Corning Costar high binding assay plates. Plates were coated overnight at RT with 10 µg/ml bait protein and thereafter blocked for 2 hours in PBS supplemented with 1% BSA before being incubated with primary antibodies in PBST (PBS + 0.2% Tween20) for 1 hour. Secondary antibodies conjugated with HRP (ThermoFisher) were diluted according to the manufacturer's instructions into PBST before incubating for 1 hour. 1-Step™ Ultra TMB-ELISA Substrate Solution (ThermoFisher Scientific) was added for 20 minutes before being supplemented with 1 M $H_2SO_4$ to stop the reaction. Colorimetric signal was measured at 450 nm.

**Western Blotting**

Protein samples were mixed with SDS loading buffer and incubated at 99°C for 2 minutes. 8-15 µg of protein was run on NuPAGE 4-12% Bis-Tris gels (Invitrogen) at 80-120 V using MOPS running buffer. Protein was transferred to a 0.45 µm nitrocellulose membrane and stained with ponceau to visualise total protein. Blocking was done with with 5% milk and PBST (PBS, 0.2% Tween20) for 1 hour. The membrane was incubated with a 5% milk-PBST mixture with primary antibodies at 4°C overnight. The membrane was washed with PBST for 15 minutes, three times before being incubated with a 5% milk-PBST mixture with secondary antibody for 1 hour. Finally, the membrane was washed with PBST for 15 minutes, three times before being developed using Li-COR Odyssey Infrared Imaging System. Images are analysed on ImageStudioLite.

**Mass spectrometry sample preparation**

Protein samples were generated by lysing cells in 8 M urea buffer. Equal 100 µg aliquots (as determined by Bradford protein quantification) were stored at -80$^{o}$C. Proteins were precipitated overnight in ice-cold acetone then re-suspended in 0.1 M triethylammonium bicarbonate (TEAB). Disulfide bonds were reduced by addition of 0.2 M tris(2-carboxyethyl)phosphine (incubated for 1 hour at 55 ºC) and free cysteines were capped by addition of 0.4 M iodoacetamide (incubated for 30 minutes at room temperature, protected from light). Proteins were digested with 2.5 µg of trypsin (Promega, UK) overnight at 37 ºC. Digestion was halted by acidification with 0.4 % trifluoroacetic acid and peptide samples were desalted using a C18 MacroSpin column (The Nest Group Inc, USA). Peptides were eluted with 80 % acetonitrile and divided to approx. 25 µg and 75 µg amounts. Both peptide solutions were dried in a vacuum centrifuge and stored at -80 ºC until required. Nine samples (biological triplicates of three conditions) of dried peptides (25 µg per sample) were each solubilised in 0.05 M TEAB and labelled with one of nine TMT mass tagging reagents (Thermo Scientific) from a 0.2 mg TMT 10-plex kit. The tenth channel was used to label a pooled sample. TMT labels were removed from the freezer and thawed at room temperature for 20 minutes. Labels were solubilised (20 µl of anhydrous acetonitrile) then one sample was combined with one label. Labels are amine-reactive and modify the sample's lysine residues and peptide N-termini (one hour incubation at room temperature). Peptide labelling efficiency was assessed by analysing an aliquot using a QExactive orbitrap (Thermo Scientific) mass spectrometer. All samples displayed > 98 % labelling efficiency so were immediately quenched (addition of 1 µl of 5 % hydroxylamine) to sequester unreacted excess label. Aliquots (5 µl) were removed from each quenched sample, combined in an Eppendorf tube and mixed thoroughly. After appropriate dilution, this mix was analysed using a QExactive orbitrap mass spectrometer to determine how much of each sample had to be combined to produce a multiplexed sample containing equal peptide amounts. A final multiplexed (10-plex) sample was produced and dried in a vacuum centrifuge. The 10-plex sample was desalted using a C18 MacroSpin column, peptides were eluted with 80 % acetonitrile and the sample was dried in a vacuum centrifuge. To improve peptide coverage and quantification depth, high pH reversed-phase fractionation column was employed. The dried sample was solubilised in 0.1 % trifluoroacetic acid, loaded onto the column (Pierce, UK) and eluted into nine separate fractions using nine elution solutions of increasing acetonitrile percentage (initial fraction 5 %; final fraction 50 % acetonitrile). The elution solution contained triethylamine to maintain a high pH value. The nine eluted fractions were dried in a vacuum centrifuge.

**Mass spectrometry**

Dried peptide fractions were re-suspended in 20 µl of 0.1% trifluoroacetic acid, sonicated for 10 minutes and centrifuged at 14,000 rpm for 3 minutes at 25 °C. Each supernatant was transferred to separate glass auto-sampler vials. Single 10 µl injections were analysed using a Fusion Lumos Tribrid orbitrap mass spectrometer coupled to an UltiMate 3000 HPLC system for on-line liquid chromatographic separation. All mass spectrometry and liquid chromatography components were sourced from Thermo Scientific, USA. The sample was loaded onto a C18 trap column (Acclaim PepMap 100; 75 µM × 2 cm) then transferred onto a C18 reversed-phase column (PepMap RSLC; 50 cm length, 75 µm inner diameter). Peptides were eluted with a linear gradient of 2–25 % buffer B (75 % acetonitrile, 20 % water, 0.1 % formic acid, 5 % DMSO) at a flow rate of 275 nl/min over 135 minutes followed by a second linear gradient of 25-40 % buffer B over 25 minutes. MS1 instrument settings: spectra were acquired in the orbitrap at 120,000 resolution with scan range of 350–1500 m/z; maximum injection time 50 milliseconds; AGC target value 4E5; 30 % RF lens setting; 90 second dynamic exclusion. MS2 instrument settings: precursor ions with charge states $z = 2–7^{+}$ were selected for MS/MS higher-energy collision-induced dissociation (HCD) fragmentation; 38 % energy; acquired in the orbitrap at 60,000 resolution; maximum injection time 105 milliseconds; AGC target value 1E5.

**Mass spectrometry data analysis**

All data were analysed with MaxQuant software version 1.6.0.13***(41)***. Default TMT 10-plex quantification and identification settings were used. MS/MS spectra were searched against a UniProtKB *Sulfolobus acidocaldarius* protein database (downloaded January 2019 containing 2,222 sequences composed of 349 Swiss-Prot reviewed and 1,873 TrEMBL un-reviewed sequences) using the Andromeda search engine. Data were also searched against a reversed decoy of the protein database and a list of common protein contaminants. MaxQuant settings: trypsin enzyme (C-terminal cleavage of arginine and lysine residues); two missed cleavages; Carbamidomethylation of cysteine as a fixed modification; N-terminal protein acetylation and methionine oxidation as variable modifications. All other MaxQuant parameters were maintained at default settings including a protein and peptide false discovery rate of 1 %. The MaxQuant output proteinGroups.txt file was opened in Perseus version 1.4.0.2(*42*). Proteins were filtered to remove proteins identified in the common contaminant list and reverse decoy database. Intensity values were log 2 transformed and filtered to include only those proteins that provided intensity values in all TMT channels. Column then row data were normalised by median subtraction. Profile, volcano and scatter plots were generated for data visualisation and interpretation.

**MCB9463 - TMT 6-plex analysis of bortezomib / wash out experiment.**

Protein samples were generated by lysing cells in 8 M urea buffer. Equal 50 µg aliquots (Bradford quantification) were submitted to the Proteomics facility. Samples were tryptically-digested for 3 hours using the iST Sample Preparation Kit (PreOmics GmbH, Germany) following the manufacturer's instructions. Eluted peptides were dried in a vacuum centrifuge and stored at -80 ºC. Six samples (biological triplicates of two conditions) of dried peptides (50 µg per sample) were solubilised in 0.05 M TEAB and labelled with one of six TMT mass tagging reagents (Thermo Scientific) from a 0.8 mg TMT 6-plex kit as described above for the TMT 10-plex kit. Labelling efficiency assessment (> 98 %), a mixing check, production of a final multiplexed (6-plex) sample, desalting and high pH fractionation were performed as described above. MS and Proteomic data analysis were also performed as described above.

**Antibody generation**

Protein polyclonal antibodies against CdvB1 were raised in chicken by Innovagen (Lund, Sweden) according to their standard protocol including IgY purification. The protein used for immunization, was expressed and purified from E. coli Rosetta (DE3) using Ni-affinity chromatography described in Methods. Peptide antibodies against the CdvB2 peptide $NH_2$-CADVNDFLRNWG-$CONH_2$ were raised in guinea pig by Innovagen (Lund, Sweden) according to their standard protocol including IgG purification.

**Immunofluorescence labelling**

Cells were fixed by stepwise addition of $4^oC$ absolute ethanol until the concentration reached 70% from a string point of 30%. Samples were washed and rehydrated in PBS-TA (PBS supplemented with 0.2% Tween20 and 1% bovine serum albumin) before incubation overnight at $25^oC$ with primary antibodies. Conjugated secondary antibodies were used for detection of target proteins (Table S3). The S-layer was stained by incubating the rehydrated sample with 200 µg/ml Concanavalin A conjugated to Alexa Fluor 647 for 1 hour. DNA was visualized by the addition of 1 µg/ml Hoechst to samples after final washes. EdU was labelled with Click-IT chemistry according to kit instructions (ThermoFisher, C10639).

**Microscopy**

Microscopy samples were prepared by coating LabTek chambered with 2% polyethyleneimine (PEI) for 30 minutes at $30^oC$. Slides were then washed with distilled water and 200 µl of stained cell suspension was added and spun for 30 minutes at 1000 RCF in a swing-bucket rotor. Super-resolution was achieved using the NanoJ liveSRRF package for Fiji(*32*, *43*). Images were captured for 1 second in 100 x 10 µs intervals on a Nikon Ti2, 1.49NA, 100x objective (Nikon CFI Apochromat TIRF 100XC oil objective) with a Photometrics Prime 95B sCMOS camera. Illumination was provided with a CoolLED pE-4000 LED illuminator. Analysis of ring diameters and nucleoid segregation was done with the ObjectJ plugin for Fiji (sils.fnwi.uva.nl/bcb/objectj/). Data was gathered from 3 biological replicates.

**Flow cytometry**

Flow cytometry analysis was carried out on BD-Biosciences LSR II and BD-Biosciences Fortessa with immunostained cells going through the following lasers and filters: lasers: 355nm, 488nm, 561nm 633nm filters: 450/50UV, 525/50 Blue, 582/15 YG, 710/50 Red. Side Scatter and Forward scatter was also recorded. Analysis was carried out on FlowJo software 10.

**Protein cloning, expression, purification for crystallography**

CdvB, CdvB1 and CdvB2 genes were amplified from genomic *S. acidocaldarius* DNA with Phusion polymerase (NEB) according to the manufacturer's instructions. The primers used in this study can be found in Table S5 and carried homologous regions to the pSUMO-YHRC plasmid (Addgene 54336). For plasmid construction the Gibson Assembly (NEB) was used according to the manufacturer's instructions. A 2000 ml culture of E. coli Rosetta (DE3) transformed with pSUMO_S1373 (CdvB), pSUMO_S0451 (CdvB1) or pSUMO_1416 (CdvB2) was cultivated in LB media supplemented with 0.1% D-glucose, 50 µg/ml Kanamycin and 34 µg/ml Chloramphenicol at 37$^{o}$C until $OD_{600}$ reached 1.5. Following 30 minutes incubation at 4$^{o}$C expression was induced with the addition of 0.5 mM IPTG overnight. Cells were harvested and resuspended in Buffer A (50 mM Tris pH 7.5, 500 mM NaCl, 5% Glycerol) supplemented with 20 mM imidazole before being lysed on ice through sonication. Lysate was heat-inactivated at 70$^{o}$C for 20 minutes before debris was pelleted at 20,000 RCF for 45 minutes. The supernatant was loaded onto an equilibrated His GraviTrap (GE Healthcare). The loaded column was washed with 15 ml Buffer A supplemented with 20 mM imidazole before being eluted with Buffer A supplemented with 250 mM imidazole. Fractions were loaded onto a PD10 Salt Exchange Column (GE Healthcare) equilibrated with Buffer A. Protein bearing fractions were supplemented with Ulp1 and left at RT overnight to cleave the sumo-tag. Samples were added to a His GraviTrap equilibrated with Buffer A, collecting elute. Aliquots were snap-frozen in liquid nitrogen and stored at -80$^{o}$C.

**Crystallization of the Saci0613/Saci0662ΔN 20S core proteasome**

The inactive Saci0613/Saci0662ΔN proteasome was overexpressed in *E. coli* and purified via heat treatment at 70°C in CR buffer (100 mM Tris-HCl pH 8, 300 mM NaCl, 10% glycerol). Stable proteasome complexes were retrieved from the supernatant using Nickel-agarose IMAC chromatography and eluted in CR buffer supplemented with 500 mM imidazole. The complex was further purified by two subsequent rounds of size exclusion chromatography in GF buffer (150 mM NaCl, 20 mM Tris pH 8.0, 5% glycerol) using a Superdex 200 10/300 GL gel filtration column. Elution fractions corresponding to fully formed proteasomes (four stacks of homo-heptameric toroids) was then concentrated to 10 mg/ml. Crystals were grown using equal volumes of CZ buffer (25% PEG 1500, 0.1M MMT buffer pH5 [200 mM DL-malic acid, 400 mM MES, 400 mM Trizma]) and 10 mg/ml Saci0613/Saci0662ΔN complex. The initial 0.6 ml drops were equilibrated against a 70 ml well and the crystals generated were then used to seed 2 ml drops equilibrated against a 60 ml well, to increase the size and resolution of the crystals. Crystals were transferred to a drop of CZ solution supplemented with 30% glycerol and immediately snap frozen in liquid nitrogen. The 3.7 Å dataset was collected from a single crystal at SOLEIL (proxima 2 beamline) France at a wavelength of 0.9801 nm. The data were integrated using XDS(*44*). The space group symmetry was assigned in POINTLESS and intensities were scales in AIMLESS(*45*). Molecular replacement and automated refinement were carried out in PHENIX(*46*–*48*). Manual refinement was carried out using COOT(*49*).

**Modelling of the Saci_0909ΔN catalytically active β-subunit**

A homology model of the active Saci_0909ΔN β subunit was generated with Modeler 9.20**(*20*)** using the crystallographically determined structure of the inactive Saci_0662ΔN β subunit. To position the catalytic threonine correctly the structures of *Archaeoglobus fulgidus* (PDB ID code 1J2Q) and *Saccharomyces cerevisiae* (PDB ID code 4NNN) were also incorporated as templates. The final model was determined by its discrete optimised protein energy (DOPE) score.

**Docking of MG132 and bortezomib**

A docking model for bortezomib was generated in CovDock(*23*) using only the Saci_0909ΔN/Saci_0662ΔN pair of subunits that form the substrate binding pocket and receptor. The catalytic threonine of Saci_0909ΔN was identified for covalent attachment to the boronic acid moiety of the inhibitor(*13*, *23*). Final models were chosen based on a combination of docking energy scores (kcal/mol) prior to the formation of the covalent bond and visual inspection to check for correspondence with the experimentally determined binding state of bortezomib in the *Saccharomyces cerevisiae* 20S proteasome (PDB ID: 2F16 and 4QVW)(*24*, *25*).

***In vitro* 20S proteasome processing assays (with and without bortezomib inhibition).**

The 20S proteasome core proteasome degradation reactions were performed as described previously(*18*). 10 µg of the GFP, or 3xUrm1-GFP, was incubated with 30 µg 20S proteasome complex in reaction buffer (20 mM Tris acetate (pH 8.0), 100 mM NaCl, 5 mM $MnCl_2$, 10 mM MgOAc, 50 mM KOAc, 1 mM $Zn(OAc)_2$, 5% glycerol and 0.02% β-mercaptoethanol) in a 75 µl reaction volume at 63°C for 0, 60, 90, or 120 min, respectively. Bortezomib (solubilized in DMSO as a 100 mM stock) was added to the appropriate reactions at a final concentration of 50 µM. After addition of 2X laemmli protein loading buffer, the products were then separated by SDS-PAGE, and visualized with Coomassie stain.

**Coarse-grained molecular dynamics simulations**

Following the notation from Yuan et al.(*50*), we modelled a fluid, spherical membrane consisting of 48,000 particles with parameters εbead−bead=4.34 kBT, ξ=4, µ=3, and rcut=1.12 σ, where σ is the membrane particle diameter which roughly corresponds to 10 nm in our simulations. The ESCRT-III filament is built of three-beaded rigid subunits which are connected by strong harmonic bonds to form a helix (bond strength K=600 kBT).

For simplicity the filament length N(t) is defined as the number of subunits that is still integrated in the filament at time t. Attraction between the ESCRT-III helix and the cell membrane is modelled by a Lennard-Jones potential

$$E_{ij} = 4\varepsilon \left((\sigma/r_{ij})^{12} - (\sigma/r_{ij})^{6}\right),\ r<r_c$$

with ε=4 kBT, σ=10 nm, rc=11.2 nm. It acts between the green beads of the ESCRT-III-filament and a membrane particle of distance $r_{ij}$.

In order to investigate how the constriction and disassembly of the ESCRT-III-filament affects the cell membrane dynamics, we ran simulations with the Molecular Dynamics package LAMMPS (https://lammps.sandia.gov/). All particles in the system experience Brownian dynamics with friction coefficient set to unity, $\gamma=m/t_0$, where m is the particle mass (set to unity for all particles) and $t_0$ is the simulation unit of time. The system is first equilibrated for $2\times10^5$ time-steps (with a fixed time-step of 0.01 $t_0$) to ensure that the ESCRT-III-filament attaches to the membrane. When the last subunit of the ESCRT-III-filament is disassembled, the simulation relaxes for another $10^5$ time-steps. The simulation results are then visualized by the open-source software OVITO(*51*). In order to compare the simulation results to the experimental ones, we measure the cell diameter d(t) at the constriction zone as a function of time. For that, we collect all membrane particles that are located in a cuboid which is fixed at the centre of the simulation box and has a width of 1 σ. The collected coordinates are then projected onto a plane and the resulting data is fitted by a circle using the Taubin method implemented in C++ by N. Chernov(*52*). Please note that we measure the cell diameter, while the ESCRT-III-ring diameter was measured in experiments. Next, we sample 108 random numbers according to the function d(t)/d0, sort them by size in descending order, rescale the values to the experimentally measured range, and take into account that the initial diameter d0 is Gaussian distributed with mean=1.26 µm and standard deviation=0.14 µm.

**Statistical analysis**
All statistical analysis and data processing was done using Microsoft Excel and GraphPad Prism 6 software. Significance was defined as $p < 0.05$. Significance levels used were *P < 0.05, **P < 0.01, ***P < 0.001 and ****P < 0.0001.

Supplementary Figure 1 Electron density map of the Saci_0613/Saci_0662ΔN proteasome

A

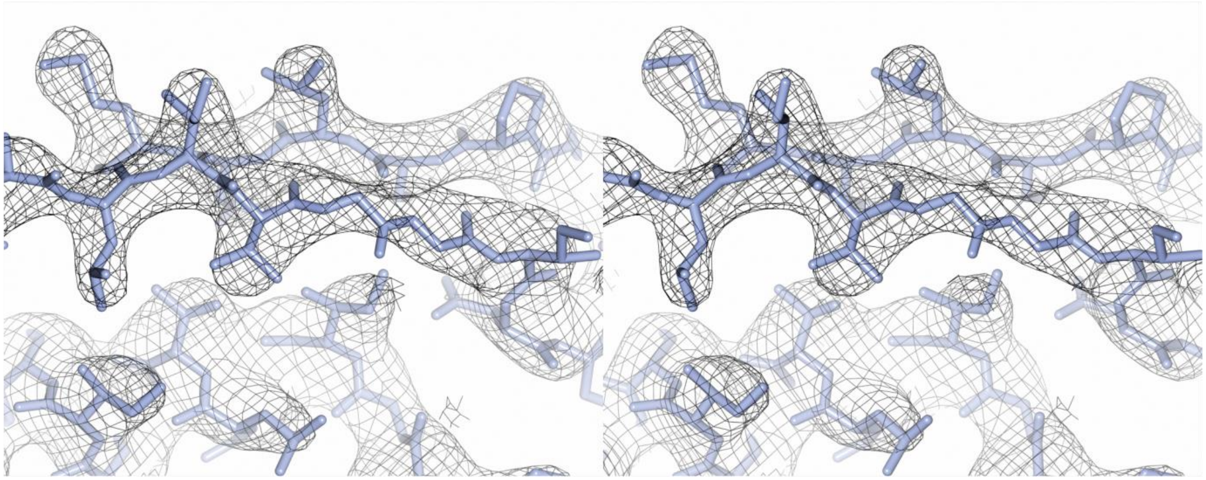

B

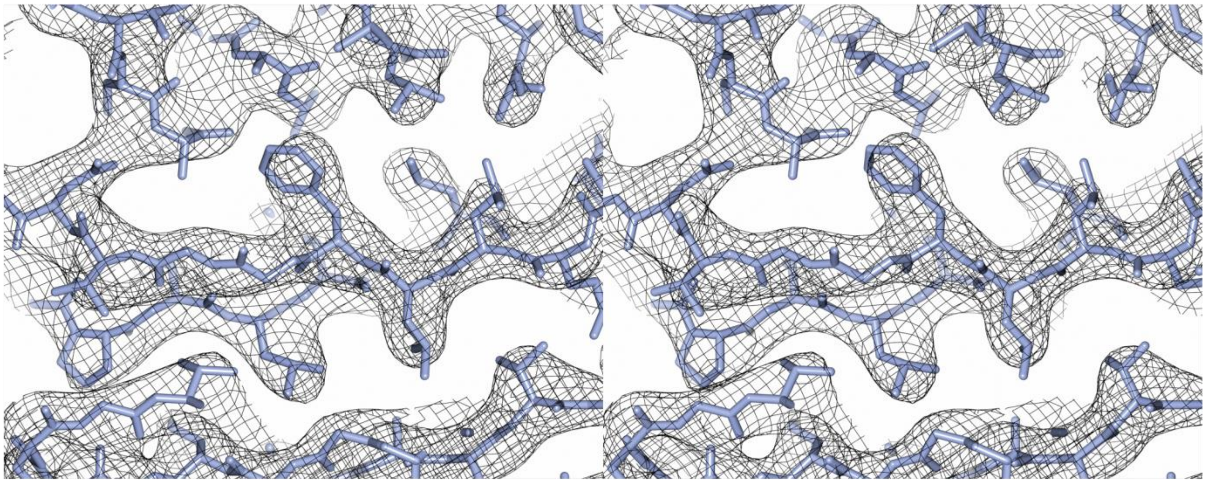

C

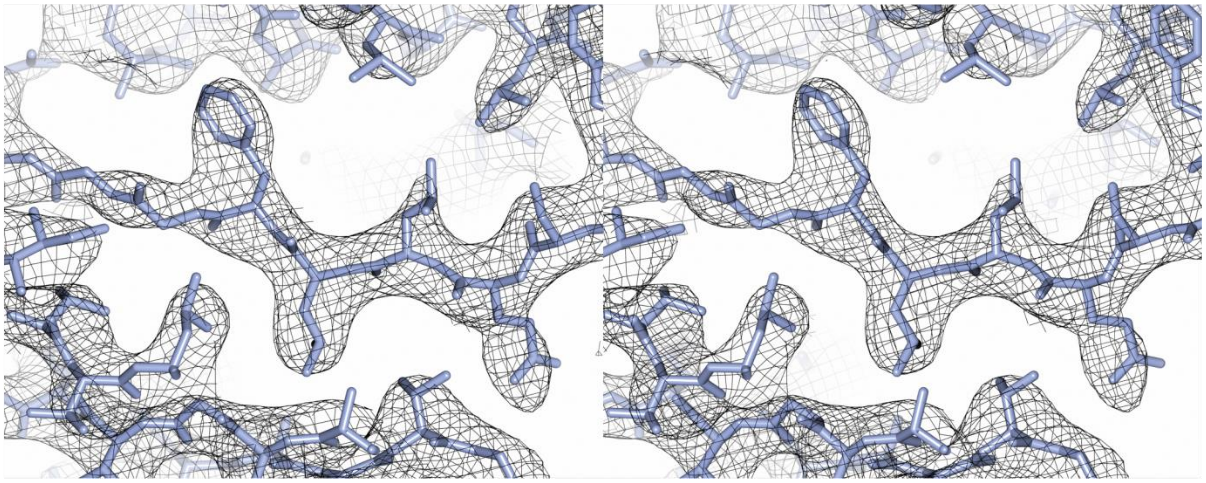

**Supplementary Figure 1** Electron density map of the Saci_0613/Saci_0662ΔN proteasome. Stereo image of a portion of the electron density map of the *S. acidocaldarius* Saci_0613/Saci_0662ΔN proteasome structure [Type of map: mtz, Contour level: 1 sigma]. (**A**) Segments from subunit B secondary structure element α-3. (**B**) subunit B secondary structure element β-5. (**C**) subunit a secondary structure element β-6.

Supplementary Figure 2 Proteasome open pore and catalytic site

**A**

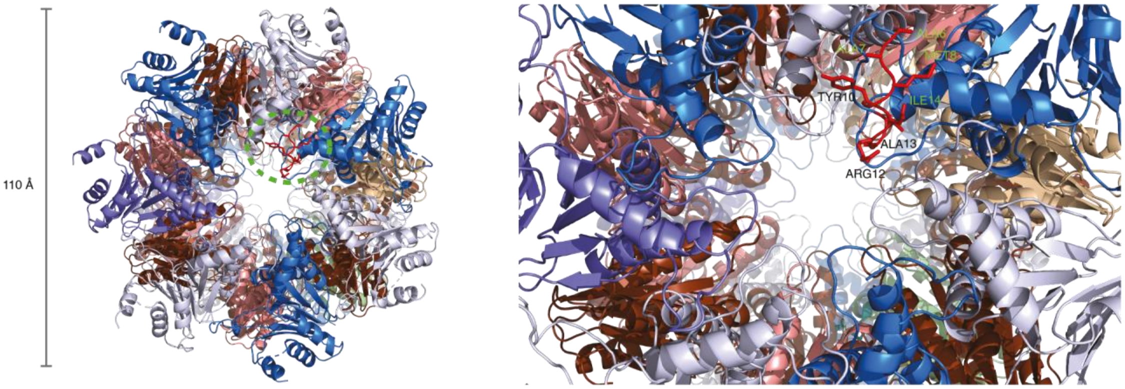

**B**

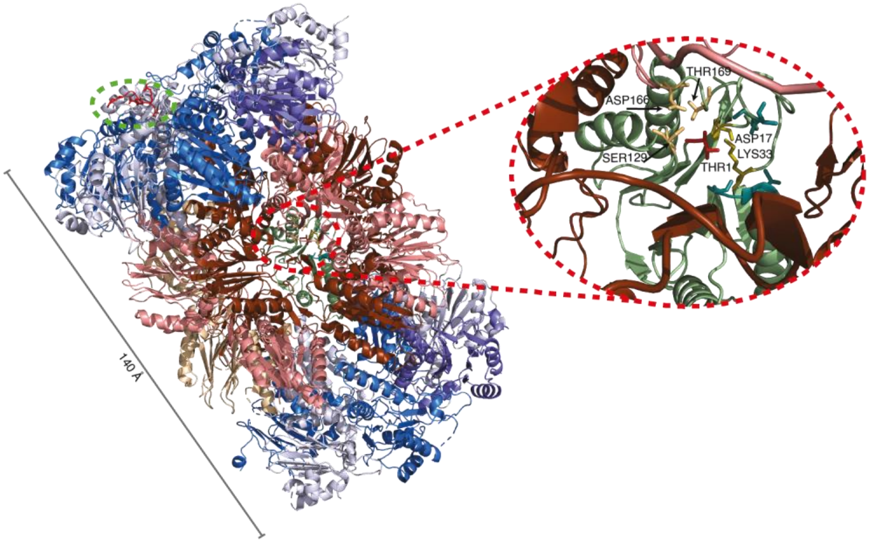

**Supplementary Figure 2** Proteasome open pore and catalytic site **(A)** (left) Top-down view of the *S. acidocaldarius* twenty-eight subunit Saci_0613/Saci_0662ΔN 20S proteasome assembly, including the modelled Saci_909ΔN active subunit, highlighting the 'open' conformation of the a-subunit gating. Saci_0613 α-subunits are coloured alternatively dark- and light-blue. Saci_0662ΔN β-subunits are coloured alternatively dark- and light-red. The extended region of α-subunit-1 residues 6-14 (AAMGYDRAI) (highlighted by the green-dashed circle) is coloured red and the side chains of these residues are shown as sticks. (right) magnified view of the pore region, highlighting the additional residues 6-14 (AAMGYDRAI) that could be built into the density map at the N-terminus of α-subunit-1. **(B)** (left) Side view of the 20S proteasome assembly with the extended region of α-subunit-1 residues 6-14 (AAMGYDRAI) highlighted by the green-dashed circle. The modelled Saci_909ΔN active subunit (green) is on the opposing side of the b-ring. (right) Magnified view through the Saci_0662ΔN b-ring, revealing the modelled Saci_909ΔN active subunit (green) and the key catalytic residues (Thr1, Asp17 and Lys33) at the active site, with three further conserved residues (Ser129, Asp166 and Thr169).

Supplementary Figure 3 Cell division cycle responses to proteasomal inhibition

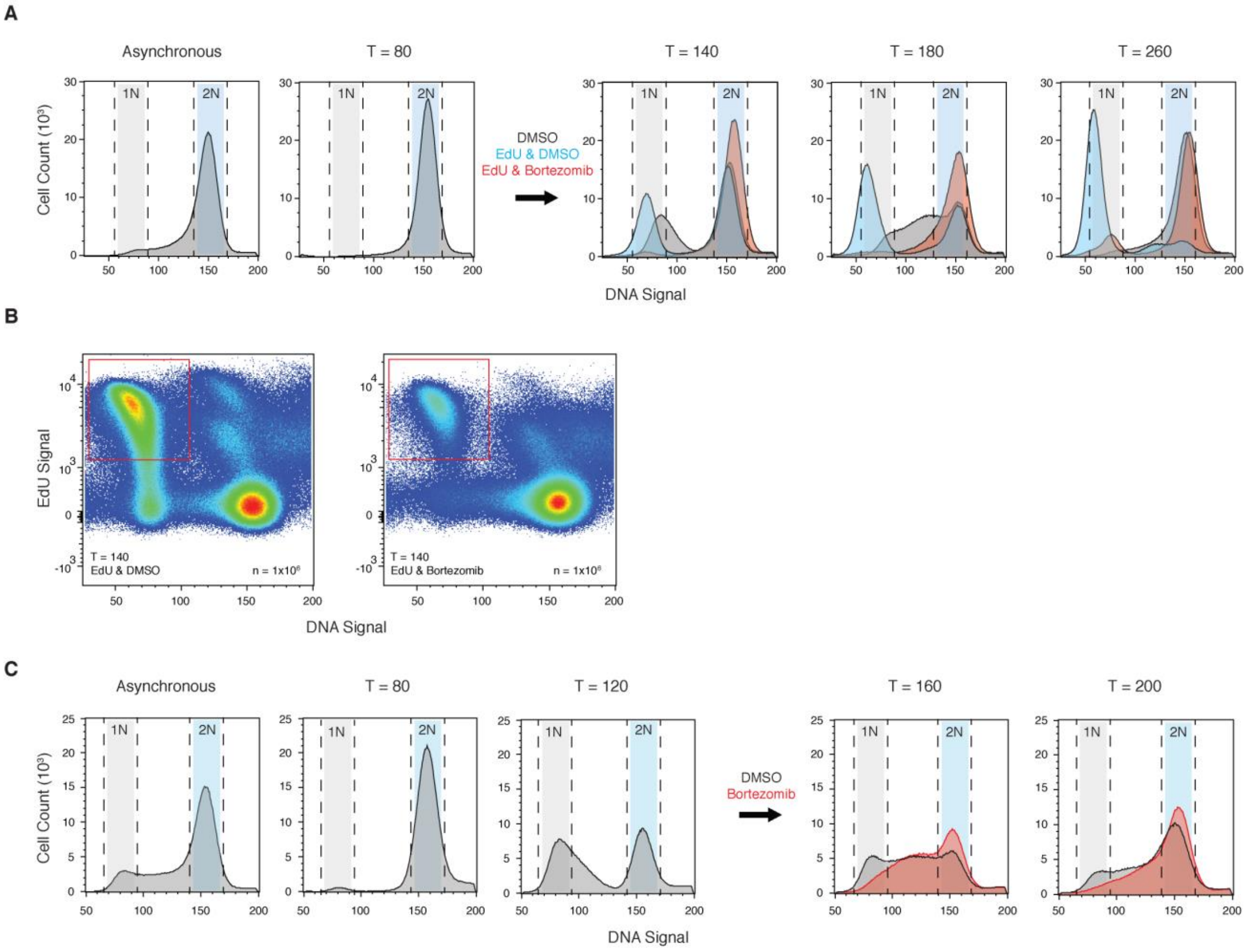

**Supplementary Figure 3** Proteasomal inhibition leads to cell division arrest. (**A**) Distribution of cells with 1N and 2N DNA content in a STK cell line population following acetic acid synchronisation and bortezomib treatment 100 min after release (pre-division). EdU was added to the synchronised population 80 min after release from acetic acid. N=3, $n=10^6$. Representative histograms shown. (**B**) Scatterplots of EdU content vs DNA content vs cell count (color code: high density red and low density blue) of a synchronised STK population 140 minutes after release, treated with DMSO or bortezomib respectively 100 min after release. The red quadrant shows the S-phase cells with EdU incorporated. (**C**) Distribution of cells with 1N and 2N DNA content in a population following acetic acid synchronisation and bortezomib treatment 120 min after release (post-division). N=3, $n=10^6$.

### Supplementary Figure 4 Proteasomal inhibition with MG132

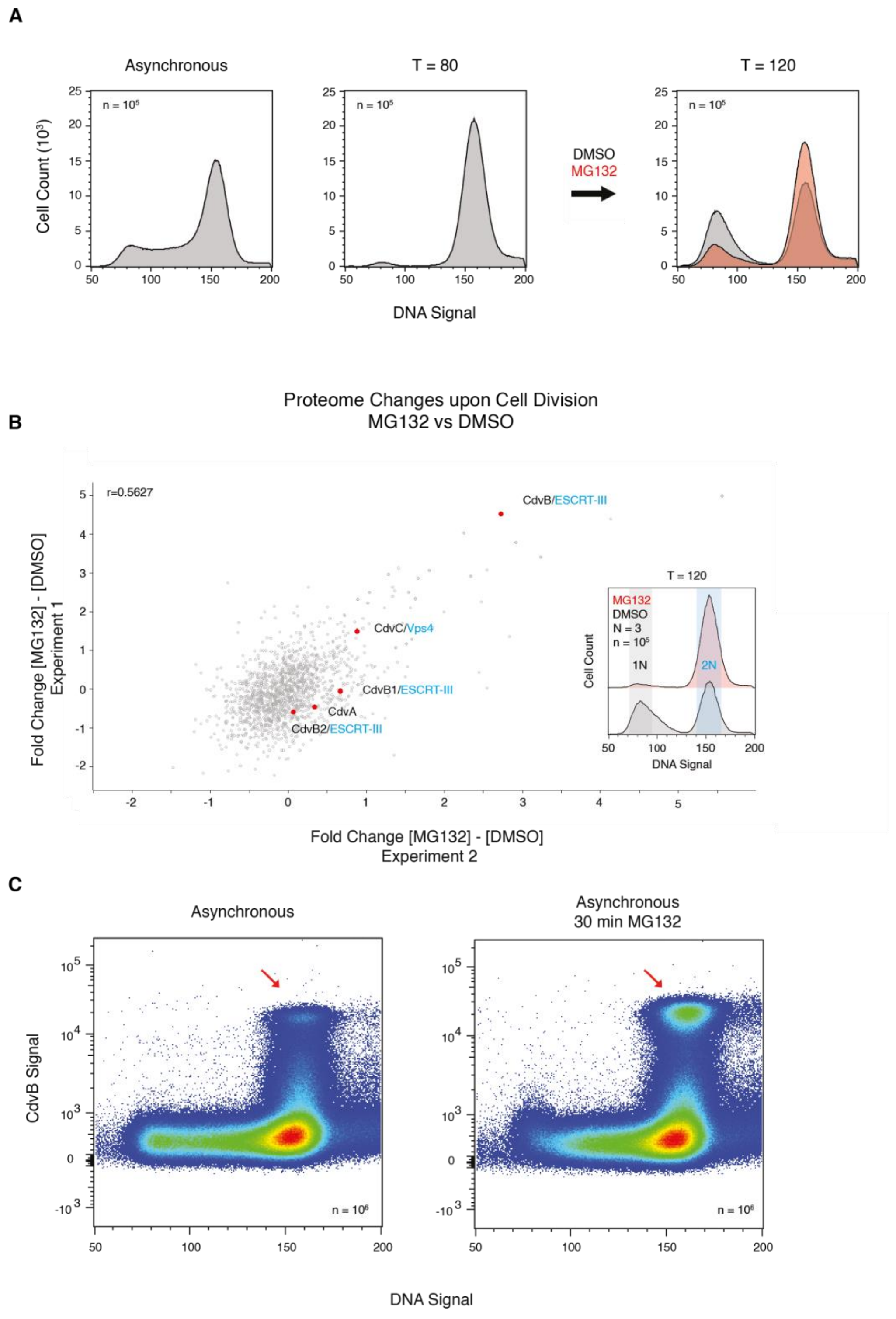

**Supplementary Figure 4** Proteasomal inhibition with MG132. (**A**) Distribution of cells with 1N and 2N DNA content in a population following acetic acid synchronisation and MG132 or DMSO treatment 80 min after release (pre-division). Representative histograms shown. N=3, n=$10^5$. (**B**) Mass spectrometry scatterplot correlating two independent replicates showing the difference in proteomes of pre-division synchronised cells treated with MG132 (1) or DMSO (2) for 40 min. Protein values were measured as the ratio of TMT intensities subtracting sample (2) from sample (1). The insert shows two representative histograms of the MG132 and DMSO treated samples. (**C**) Scatterplots of CdvB content vs DNA content vs cell count (color code: high density red and low density blue) of asynchronous populations untreated (left) and treated with MG132 (right) for 30 min. The red arrow highlights the increase in the number of cells within the population with peak CdvB signal. Representative scatterplots shown. N=3, n=$10^6$.

Supplementary Figure 5 Cdv transcript and protein level response to proteasomal inhibition

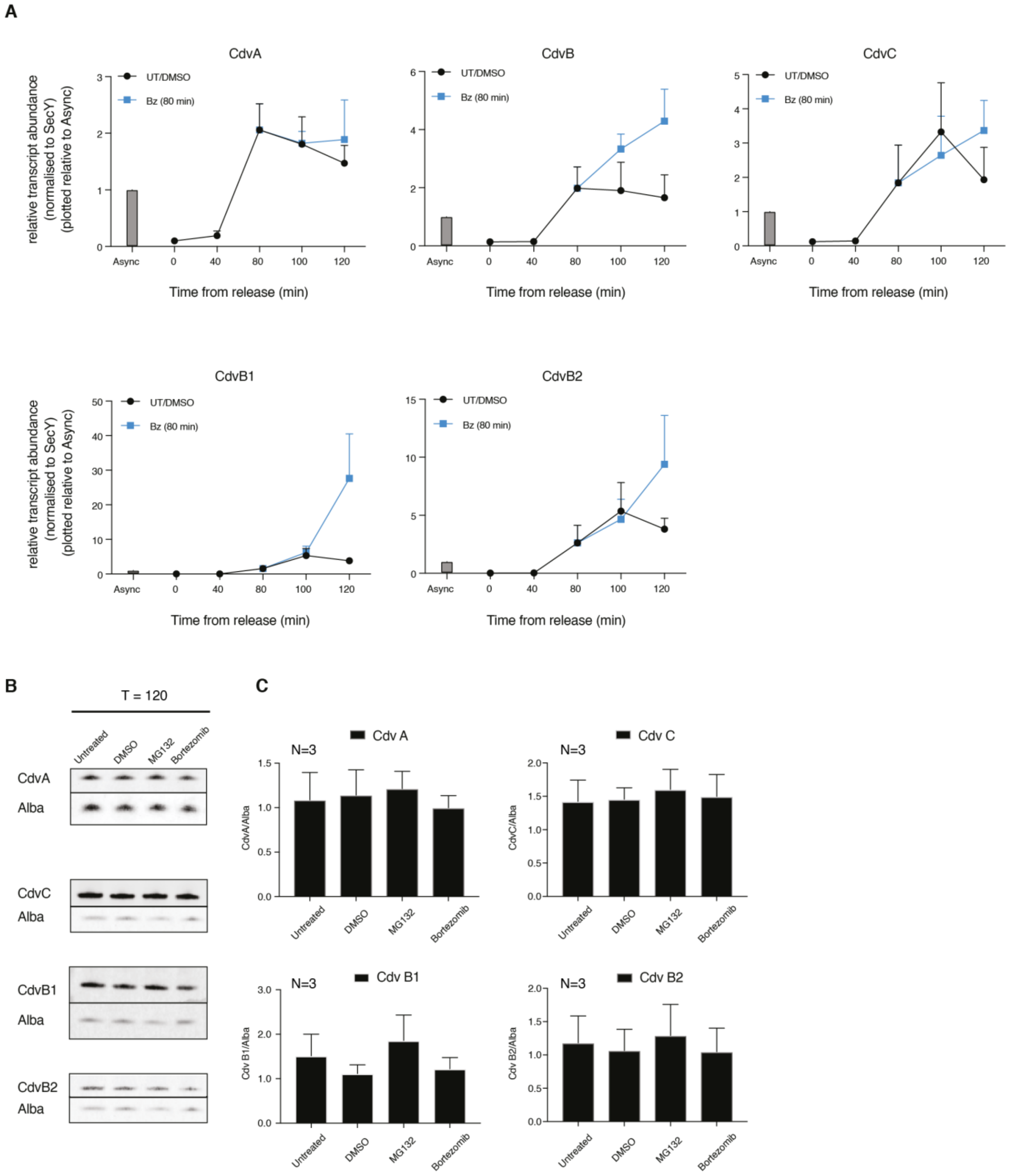

**Supplementary Figure 5** Cdv transcript and protein level response to proteasomal inhibition. (**A**) qPCR transcript abundance of Cdv proteins in over time in an acetic acid synchronised population with bortezomib or DMSO treatment 80 min after release (pre-division). (**B**) Western blots with CdvA, CdvC, CdvB1, and CdvB2 protein levels with Alba as a loading control. All samples were taken 120 min after release from acetic acid synchronisation, after having been left untreated or treated with DMSO, MG132 or bortezomib 80 minutes after release. Representative blots shown. (**C**) Quantification of the western blots. N=3.

Supplementary Figure 6 Targeted degradation of CdvB triggers constriction of CdvB1/B2

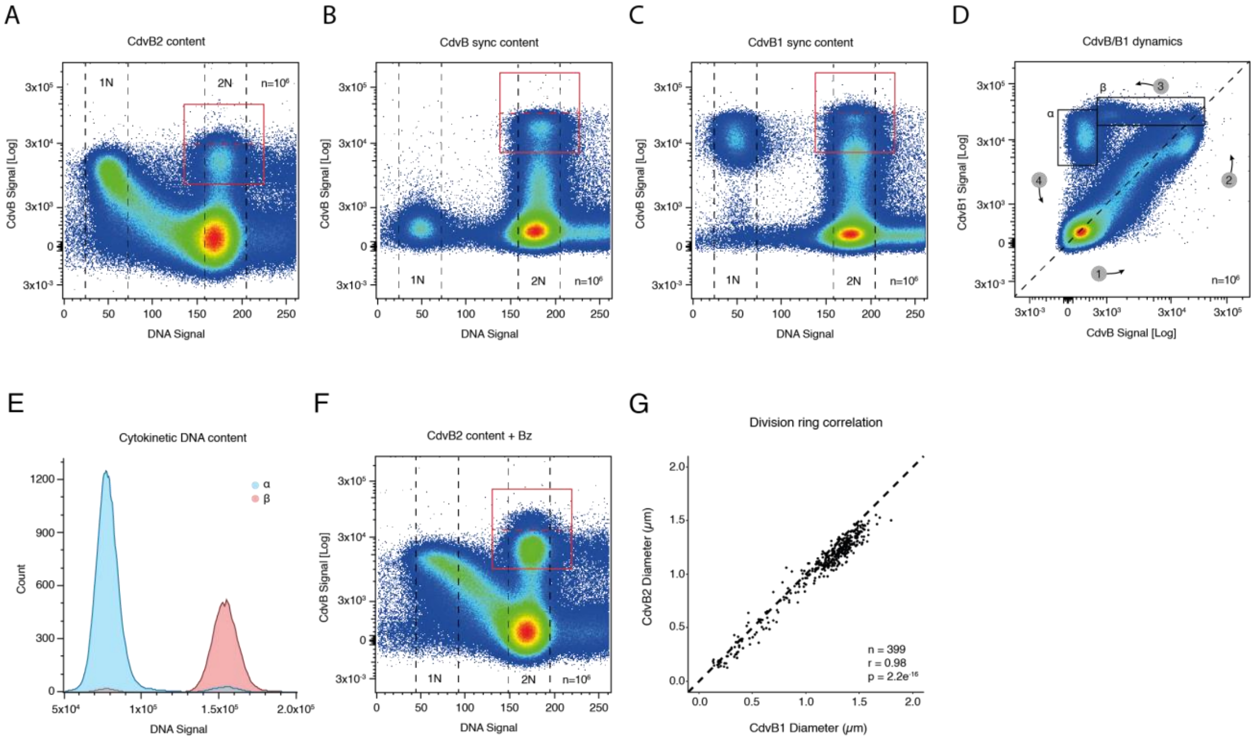

**Supplementary Figure 6** Targeted degradation of CdvB triggers constriction of CdvB1/B2 **(A)** Scatterplot of DNA versus CdvB2 content for an asynchronous culture. Representative plots shown, N = 3, n = $10^6$. **(B, C)** Scatterplot of DNA versus CdvB and CdvB1 content for a synchronous culture. Representative plots shown, N = 3, n = $10^6$. **(D)** Scatterplot of CdvB versus CdvB1 content in a synchronous culture showing the alternating accumulation and loss of protein (N = 3, n = $10^6$). **(E)** DNA distribution of cells in boxes shown in (**C**) with α in blue and β in red showing a shift from 2N to 1N. **(F)** Scatterplot of DNA versus CdvB2 content for an asynchronous culture treated with bortezomib. Representative plots shown, N = 3, n = $10^6$. **(G)** Correlative analysis of CdvB1 and CdvB2 ring diameters. All quantifications cover >6 fields of view over three biological replicates.

Supplementary Figure 7 Saci ESCRT-III antibody specificities

**A**

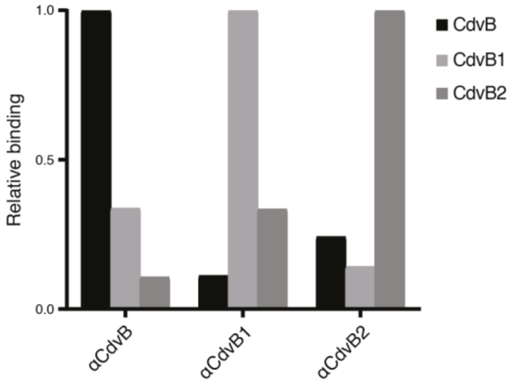

**Supplementary Figure 7 (A)** In vitro analysis of CdvB, CdvB1 and CdvB2 antibody specificities. CdvB (black), CdvB1(grey) and CdvB2 (dark grey) was coated onto a well before detection with antibodies, shown as αTarget.

Supplementary Table 1 Data Collection and refinement statistics

| | Saci_0613/Saci_0662ΔN Proteasome |
|---|---|
| **Data Collection** | |
| Space group | P 21 21 21 |
| **Cell Dimensions** | |
| a, b, c (Å) | 108.289 193.654 323.671 |
| a, b, g (°) | 90 90 90 |
| Wavelength (Å) | 0.9801 |
| Resolution range (Å) | 48.19 - 3.698 (3.83 - 3.698)* |
| Mean I/sigma(I) | 5.64 (1.56) |
| Completeness (%) | 99.21 (96.06) |
| Wilson B-factor | 82.08 |
| Multiplicity | 2.0 (2.0) |
| Refinement | |
| Total reflections | 144662 (13665) |
| Unique reflections | 73138 (6972) |
| R-work | 0.2873 (0.3422) |
| R-free | 0.3343 (0.3992) |
| Reflections used in refinement | 73060 (6967) |
| Reflections used for R-free | 3676 (347) |
| Number of non-hydrogen atoms | 41436 |
| macromolecules | 41436 |
| Protein residues | 5391 |
| RMS(bonds) | 0.002 |
| RMS(angles) | 0.57 |
| Ramachandran favored (%) | 94.7 |
| Ramachandran allowed (%) | 4.96 |
| Ramachandran outliers (%) | 0.34 |
| Rotamer outliers (%) | 0.61 |
| Clashscore | 11.63 |
| Average B-factor | 76.94 |

*Statistics for the highest-resolution shell are shown in parentheses.

Supplementary Table 2 Mass Spectrometry tables with Cdv proteins

**A**

| | CdvA | CdvB | CdvC | CdvB1 | CdvB2 |
|---|---|---|---|---|---|
| **Experiment 1:** [Bortezomib] - [Bortezomib Wash-out] | 0.36 | 2.53 | -0.07 | -0.44 | 0.32 |
| **Experiment 2:** [Bortezomib] - [Bortezomib Wash-out] | 0.26 | 2.84 | 0.54 | 1.03 | 0.53 |
| **Experiment 3:** [Bortezomib] - [Bortezomib Wash-out] | -0.40 | 2.44 | -0.08 | -0.14 | 0.11 |

**B**

| | CdvA | CdvB | CdvC | CdvB1 | CdvB2 |
|---|---|---|---|---|---|
| **Experiment 1:** [MG132] - [DMSO] | -0.30 | 1.44 | -1.08 | -0.26 | -0.25 |
| **Experiment 2:** [MG132] - [DMSO] | -0.45 | 4.53 | 1.51 | -0.04 | -0.58 |
| **Experiment 3:** [MG132] - [DMSO] | 0.34 | 2.72 | 0.89 | 0.67 | 0.07 |

Supplementary Table 3 List of antibody host species and secondary antibodies used.

| Target | Primary antibody host | Secondary antibody |
|---|---|---|
| CdvB | Rabbit | Goat anti-rabbit Alexa Fluor 546 |
| CdvB1 | Chicken | Goat anti-chicken Alexa Fluor 488 |
| CdvB1 | Chicken | Goat anti-chicken Alexa Fluor 555 |
| CdvB2 | Guinea pig | Goat anti-guinea pig Alexa Fluor 488 |

Supplementary Table 4 List of primers used in cloning of plasmids used for protein expression.

| Gene | Sequence |
|---|---|
| cdvB-F | GATATTATTGAGGCTCACAGAGAACAGATTGGTGGGATGTTTGATAAGTTATCGATAATTTT |
| cdvB-R | GCAGCCGGATCTCAGTGGTGGTGGTGGTGGTGCTCGAGTTAACCCTCAAGAACAATTAGAC |
| cdvB1-F | GATATTATTGAGGCTCACAGAGAACAGATTGGTGGGATGACCGCAATATATATATATTGACC |
| cdvB1-R | GCAGCCGGATCTCAGTGGTGGTGGTGGTGGTGCTCGAGTTAGCTCTTTTGTTGCTGTGTAA |
| cdvB2-F | GATATTATTGAGGCTCACAGAGAACAGATTGGTGGGATGGCAGATGTGAATGATTTTCTGAG |
| cdvB2-R | GCAGCCGGATCTCAGTGGTGGTGGTGGTGGTGCTCGAGTTATTTTTGATTTGCGTTTGTTTTTT |

Supplementary Table 5 List of primers used for RT-qPCR analysis

| Saci annotation | Gene | Sequence |
|---|---|---|
| Forward Saci_0574 | SecY | ACTCTTGCTTGACGAGATGATAC |
| Reverse Saci_0574 | SecY | ACTCTGTACGGAGACTATTCCA |
| Forward Saci_1374 | CdvA | GGACAGATGGAGAAGATAAGGAAG |
| Reverse Saci_1374 | CdvA | TGAGCAACTTGGTCCTCTATTG |
| Forward Saci_1373 | CdvB | ACTGGTGCATTAAGCGAGAA |
| Reverse Saci_1373 | CdvB | TTGGTAACTCTGAAGGTGGATG |
| Forward Saci_1372 | CdvC | GCTGTAGCTAATGAGATCGACTCATATT |
| Reverse Saci_1372 | CdvC | CAGCCTCGCCTAGCCACTTA |
| Forward Saci_0451 | CdvB1 | GGTGAACTAGGTGAAGGTTTACA |
| Reverse Saci_0451 | CdvB1 | CTGTCTTGCCTCTGGTGAATAG |
| Forward Saci_1416 | CdvB2 | CTCTTCTCCAGAGGCAAGAAAG |
| Reverse Saci_1416 | CdvB2 | CCGAAGTTGCAAATGATGGTAAA |

**Supplementary Movie 1** Constriction simulation without filament disassembly
**Supplementary Movie 2** Constriction simulation with filament disassembly
